# ARID1B controls transcriptional programs of axon projection in the human corpus callosum

**DOI:** 10.1101/2023.05.04.539362

**Authors:** Catarina Martins-Costa, Vincent A. Pham, Andrea Wiegers, Jaydeep Sidhaye, Balint Doleschall, Maria Novatchkova, Thomas Lendl, Marielle Piber, Angela Peer, Paul Möseneder, Marlene Stuempflen, Siu Yu A. Chow, Rainer Seidl, Daniela Prayer, Romana Höftberger, Gregor Kasprian, Yoshiho Ikeuchi, Nina S. Corsini, Jürgen A. Knoblich

**Affiliations:** Institute of Molecular Biotechnology of the Austrian Academy of Sciences (IMBA), Vienna BioCenter (VBC); Vienna, Austria; Vienna BioCenter PhD Program, Doctoral School of the University of Vienna and Medical University of Vienna; Vienna, Austria; Department of Biomedical Imaging and Image-guided Therapy, Medical University of Vienna; Vienna, Austria; Institute of Industrial Science, The University of Tokyo; Tokyo, Japan; Institute for AI and Beyond, The University of Tokyo; Tokyo, Japan; Department of Pediatrics and Adolescent Medicine, Medical University of Vienna; Vienna, Austria; Division of Neuropathology and Neurochemistry, Medical University of Vienna; Vienna, Austria; Department of Neurology, Medical University of Vienna, Vienna, Austria

**Keywords:** mSWI/SNF, ARID1B, SATB2, corpus callosum agenesis, cerebral cortex, neural organoids.

## Abstract

Mutations in *ARID1B*, a member of the mSWI/SNF complex, cause severe neurodevelopmental phenotypes with elusive mechanisms in humans. The most common structural abnormality in the brain of ARID1B patients is agenesis of the corpus callosum (ACC). This condition is characterized by a partial or complete absence of the corpus callosum (CC), an interhemispheric white matter tract that connects distant cortical regions. Using human neural organoids, we identify a vulnerability of callosal projection neurons (CPNs) to *ARID1B* haploinsufficiency, resulting in abnormal maturation trajectories and dysregulation of transcriptional programs of CC development. Through a novel *in vitro* model of the CC tract, we demonstrate that *ARID1B* mutations reduce the proportion of CPNs capable of forming long-range projections, leading to structural underconnectivity phenotypes. Our study uncovers new functions of the mSWI/SNF during human corticogenesis, identifying cell-autonomous defects in axonogenesis as a cause of ACC in ARID1B patients.

Graphical abstract
Human callosal projection neurons are vulnerable to *ARID1B* haploinsufficiency.
**(Top)** During healthy development, callosal projection neurons (CPNs) project long interhemispheric axons, forming the corpus callosum (CC) tract, which can be modeled *in vitro*. **(Bottom)** In ARID1B patients, transcriptional dysregulation of genetic programs of CC development reduces the formation of long-range projections from CPNs, causing CC agenesis *in vivo* and underconnectivity phenotypes *in vitro*.

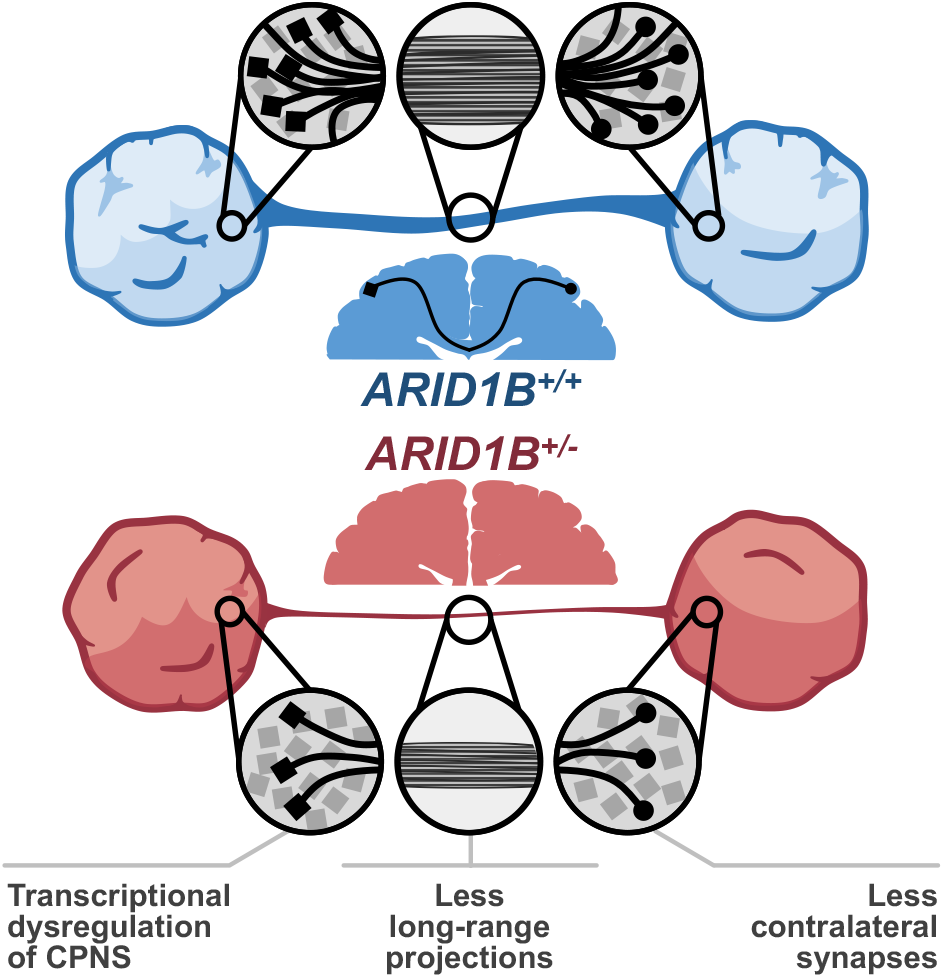

## Introduction

The mammalian SWItch/Sucrose Non-Fermentable (mSWI/SNF) machinery regulates the interconvertibility between heterochromatin and euchromatin, thereby modulating gene expression during neurodevelopment (*1*). Diverse mSWI/SNF complex configurations result from the assembly of 29 different proteins, forming cell-type- and cell-function-specific combinations of 10-15 subunits (**Fig.S1A**); these include a core ATPase, as well as DNA- and protein-binding components that modulate the targeting and activity of the complex (*1–3*). Thus, the temporal progression of mSWI/SNF subunit expression is pivotal in the epigenetic regulation of neurodevelopment, controlling processes of neural progenitor proliferation and differentiation, cell-fate determination and lineage progression, neuronal migration and maturation, as well as neurite and synapse formation (*1*). Given these ubiquitous functions, mutations in the genes encoding mSWI/SNF members are strongly linked to neurodevelopmental and cognitive dysfunction phenotypes (*1, 4, 5*).

The protein AT-rich interaction domain 1B (ARID1B, also known as BAF250B) is a core component of the mSWI/SNF complex, mediating DNA-binding in neural progenitors and neurons (**Fig.S1A**). Mutations in *ARID1B* account for 70% of cases of Coffin-Siris syndrome (*6*) and confer a heightened risk for the development of intellectual disability (ID) and autism spectrum disorder (ASD) (*5, 7, 8*). Alongside intellectual and behavioral phenotypes, ARID1B patients commonly present a striking abnormality in brain structure: agenesis of the corpus callosum (ACC) (*9, 10*).

The corpus callosum (CC) is an interhemispheric white-matter tract that connects opposing cortical regions (*11–13*). During CC development, SATB2^+^ callosal projection neurons (CPNs) respond to axon guidance cues produced at the brain midline and extend long-range callosal axons that cross to the contralateral hemisphere and establish functional synaptic connections (*14–17*) (**Fig.1A**). ACC is a neurodevelopmental condition characterized by a partial or complete absence of the CC (*18, 19*). It affects about 1 in 4,000 newborns and 3-5% of children with intellectual disability (*20, 21*), being associated with a broad spectrum of symptoms that range from mild motor incoordination to severe impairments in cognition, attention, language, and social interaction (*18, 19*). Although the aetiology of the disease is often unknown, it is estimated that 30-45% of ACC cases are caused by genetic mutations (*22, 23*).

**Figure 1.**
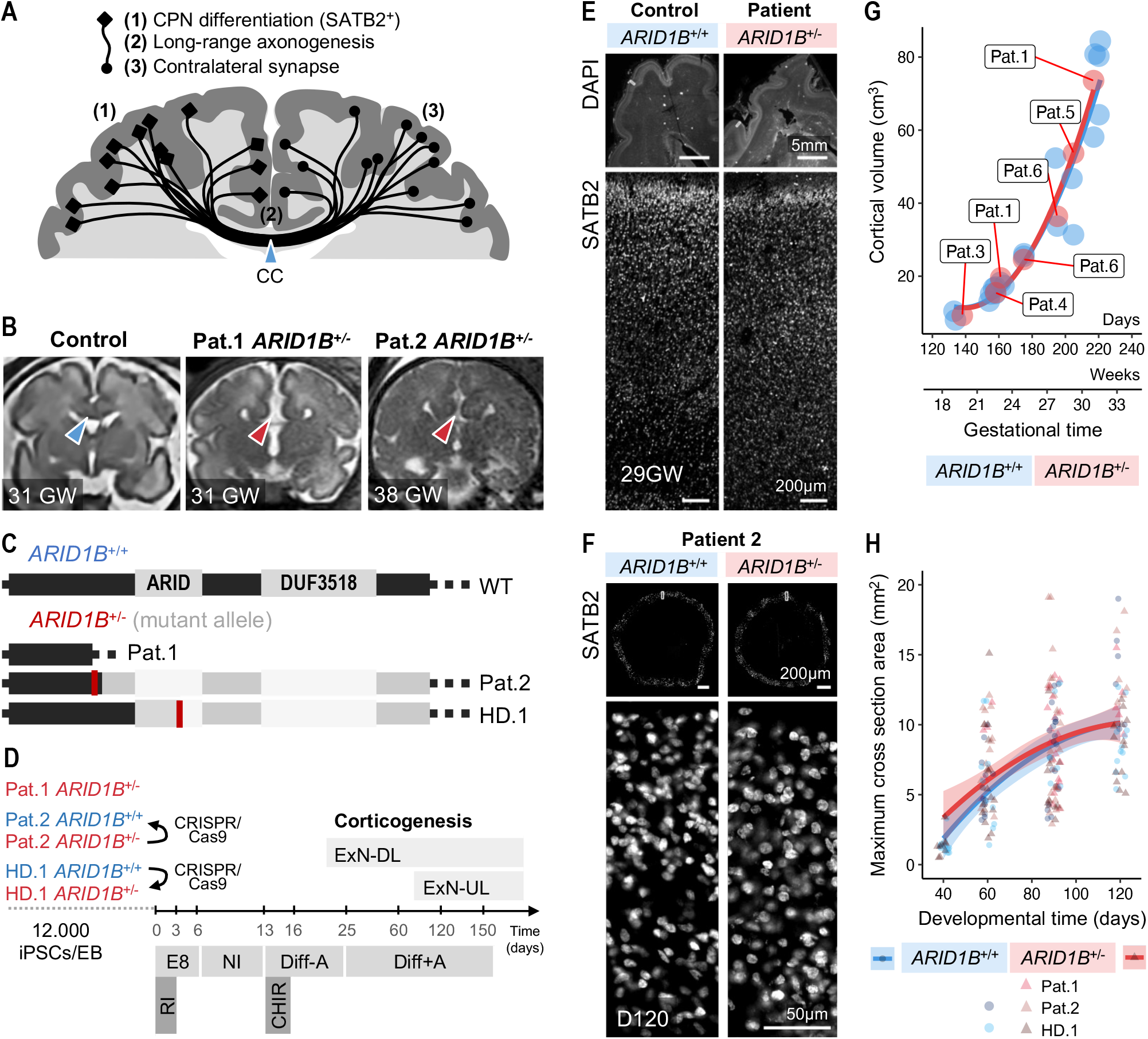
SATB2^+^ neurons are generated in *ARID1B*^+/-^ patient brains and organoids. **(A)** Schematic representation of the corpus callosum tract (CC, blue arrowhead) and sequential steps involved in its development. **(B)** Fetal MRI of a healthy control brain at 31 gestational weeks (GW), Patient 1 at 31 GW, and Patient 2 at 38 GW. Note the presence of the corpus callosum in the healthy control (blue arrowhead), and the atypical “steer horn” appearance of the frontal horns of the lateral ventricles indicating corpus callosum agenesis in the patients (red arrowheads). **(C)** Schematic representation of the wild-type *ARID1B* exon sequence, and of the mutant alleles present in Pat.1, Pat.2 and HD.1. The ARID1B protein has two functional domains: AT-rich interaction domain (ARID, spanning exons 13 and 14), which mediates DNA binding; and DUF3518, required for protein-protein interactions within the mSWI/SNF complex (*1*). Patient 1 harbors a heterozygous microdeletion that spans *ARID1B* from exon 8; Pat.2 and HD.1 *ARID1B*^+/-^ iPSCs harbor heterozygous frame-shift mutations before or on top of the ARID domain, respectively (red lines). Faded regions in Pat.2 and HD.1 represent the sequence after an early stop codon. **(D)** Protocol used for telencephalic organoid differentiation. Deep-layer excitatory neurons (ExN-DL) are generated from early stages of development; upper-layer ExNs (ExN-UL) emerge at around day 70 of development. **(E-F)** Immunostaining of SATB2 in healthy control and *ARID1B*^+/-^ patient brains at 29 GW **(E)** and in Pat.2 iPSC-derived organoids at D120 **(F)**. **(G)** Fetal MRI-based super-resolution reconstruction and growth quantification of the cortical plate volumes in healthy fetuses (blue) and six fetal *ARID1B* patients (red). Patient 1 has been included in this study. **(H)** *ARID1B*^+/+^ and *ARID1B*^+/-^ organoid growth during the generation of deep- and upper-layer neurons.

*ARID1B* mutations are one of the major causal links between genetic abnormalities and corpus callosum defects in humans. In fact, one third of ARID1B patients have ACC, and *ARID1B* mutations are present in 10% of patients with ACC in association with intellectual disability (*5, 6, 8, 9, 24, 25*). Several studies have begun to shed light onto the cognitive dysfunction phenotypes seen in patients. *ARID1B* haploinsufficiency has been shown to affect the differentiation of human neurons in 2D models systems (*26, 27*), and to lead to ID- and ASD-related phenotypes in *in vivo* mouse models (*28–31*) and *in vitro* human neural organoids (*32, 33*). However, although complete ACC is frequent in humans, this phenotype has not been recapitulated in *Arid1b* mutant mice (*28*). Also, CC formation is yet to be investigated using *in vitro* models of human neurodevelopment. Thus, the relationship between *ARID1B* haploinsufficiency and ACC is currently unclear.

Defects in neuronal specification or migration, axon outgrowth or guidance, or contralateral targeting can result in abnormal CC development (*19*). Here, we leveraged clinical data, post-mortem tissue analysis, and patient iPSC-derived neural organoids to study the impact of *ARID1B* mutations on cortical and CC development. We focused on cell-autonomous processes of CPN biology, namely fate-acquisition, transcriptional signatures, axon outgrowth and fasciculation, and synapse formation. Overall, we identified a crucial effect of ARID1B on transcriptional regulation and projection development in CPNs, whose defects suggested a novel cause of ACC in ARID1B patients.

## Results

To study how *ARID1B* mutations influence cortical development, we derived iPSCs from two patients diagnosed with CC malformations by fetal magnetic resonance imaging (MRI) (**Fig.1B**, **Fig.S1B**). Patient 1 (Pat.1 *ARID1B*^+/-^) had a short and truncated CC (partial ACC); genetic analysis identified 6q25 microdeletion syndrome (*10, 34–36*) encompassing part of *ARID1B* (from exon 8) and two downstream genes (**Fig.1C**). Patient 2 (Pat.2 *ARID1B*^+/-^) had complete ACC and harbored a heterozygous single-nucleotide duplication that led to an early stop codon upstream of the functional domains of ARID1B (**Fig.1C**); isogenic control cell lines were generated by scarless CRISPR-based genome editing (Pat.2 *ARID1B*^+/+^). Both patients presented features of Coffin-Siris syndrome (*37*), including intellectual disability. To analyze the effects of *ARID1B* mutations in a healthy control genetic background, we used iPSCs derived from a healthy donor (HD.1 *ARID1B*^+/+^) and introduced a heterozygous frame-shift mutation in the ARID domain, which resulted in an early stop codon (HD.1 *ARID1B*^+/-^) (**Fig.1C**). All experiments were performed with three *ARID1B*^+/-^ cell lines and two *ARID1B*^+/+^ isogenic controls (**Fig.1C**, **Fig.S2A-B**).

### Reduced levels of ARID1B protein do not impair iPSC maintenance or telencephalic differentiation

To analyze the effects of the identified *ARID1B* mutations on ARID1B protein abundance, we performed western blot (WB). *ARID1B*^+/-^ iPSCs showed a two-fold reduction of ARID1B protein levels compared to *ARID1B*^+/+^ isogenic controls (**Fig.S2C**). This suggests that Pat.2 and HD.1 mutations lead to nonsense-mediated mRNA decay due to premature termination of translation, as seen in most ARID1B patients (*3*). To determine if a reduction in ARID1B protein levels affected the pluripotent state of iPSCs, we assessed the expression of pluripotency markers SSEA-4 and TRA-1-60 by immunostaining and flow cytometry. The latter showed that over 94% of *ARID1B*^+/-^ iPSCs expressed both markers (**Fig.S2D**), indicating that reduced ARID1B levels did not affect pluripotency, as previously described (*27*). However, in 2D models of neural differentiation, *ARID1B* haploinsufficiency has been shown to impair exit from pluripotency during neural crest formation (*27*), and to promote premature differentiation of NPCs to neurons (*26*). To study how ARID1B mutations affect cortical specification in a 3D system, we generated telencephalic organoids enriched in dorsal-cortical tissue from *ARID1B*^+/-^ iPSCs and isogenic controls (*38*) (**Fig.1D**). Organoid formation was not impaired by *ARID1B* mutations and, at early developmental stages (day 60, D60), organoids presented neural rosettes with stratified organization of SOX2^+^ radial glia (RGs), TBR2^+^ dorsal intermediate progenitor cells (IPCs), and CTIP2^+^ early-born excitatory neurons (**Fig.S3A**), indicative of successful dorsal cortical specification. To probe the levels of ARID1B and other mSWI/SNF complex member proteins at this stage of differentiation, we performed WB and immunostaining in D60 organoids. Both in neural progenitors and neurons, ARID1B levels remained reduced (**Fig.S3B-C**), while the levels of the interchangeable mSWI/SNF core component ARID1A, and of other subunits involved in telencephalic development, BAF155 and BAF170 (*39*) (**Fig.S3B**, **Fig.S3D-F**), were unaffected. These results show that *ARID1B* haploinsufficiency did not impair iPSC maintenance or neural differentiation in the telencephalic organoid system.

### Differentiation of SATB2^+^ neurons is unaffected by ***ARID1B* mutations**

Callosal projection neurons are marked by the expression of SATB2 and born during late corticogenesis, making up most of the upper cortical layers (*16, 17, 40*). To understand how *ARID1B* mutations affect cortical development, we assessed the presence of SATB2^+^ neurons in patient cortices and organoids. SATB2^+^ neurons were abundant in *ARID1B*^+/-^ fetal cortices at gestational week (GW) 29 (**Fig.1E**). Similarly, despite the persistently lowered levels of ARID1B protein in mutant samples (**Fig.S4A**), *ARID1B*^+/+^ and *ARID1B*^+/-^ late-stage organoids (day 120, D120) showed comparable density of SATB2^+^ neurons at their outer-most surface (**Fig.1F**, **Fig.S4B-D**). In accordance with these observations, progression of fetal cortical volume measured from MRIs (**Fig.1G**, **Fig.S1C**) and of organoid size (**Fig.1H**) was comparable between patient and control groups. Together, these findings show that *ARID1B* mutations did not prevent the formation of the cerebral cortex, including the differentiation of SATB2^+^ neurons, the main population contributing axons to the CC.

### *ARID1B* mutations alter the lineage progression of excitatory cell types

Dynamic changes in the composition of the mSWI/SNF complex are developmental regulators of gene expression and, therefore, lineage progression (*7*). As such, perturbation of mSWI/SNF complex members, including ARID1B (*26*), has profound effects on neurogenic gene expression programs (*41*). To assess the impact of *ARID1B* haploinsufficiency on transcriptional programs in telencephalic organoids, we used single-cell RNA sequencing (scRNAseq) at D120 of organoid development (**Fig.2**). To account for clonal heterogeneity and batch-to-batch variability of gene expression (*42*), we performed four different sequencing experiments, resulting in five independent pairwise comparisons between organoids from different cell lines, clones, and batches (**Fig.2A**, **Fig.S5A**, details in **Materials and Methods**). By performing UMAP projection and unbiased clustering of 54.480 cells, we identified seven clusters of telencephalic populations: interneurons (INs), dividing progenitors, RGs, IPCs, and three populations of increasingly mature deep- and upper-layer excitatory neurons (ExN1, ExN2, ExN3) (**Fig.2B, Fig.S5A-B**). These clusters were characterized by the expression of known cell-type marker genes (**Fig.2C, Fig.S5C-D**). Distinct effects in cluster composition could be identified. An expansion of the ventral telencephalic lineage was observed in *ARID1B*^+/-^ organoids (**Fig.S5E**), a finding that has been previously described and is thought to contribute to the ID/ASD phenotype of the patients (*32, 33*). However, due to the guided differentiation protocol, organoids were mainly composed of dorsal cortical tissue (**Fig.S5E**). Importantly, within the excitatory lineage (from IPCs to ExN3, **Fig.S5F**), *ARID1B*^+/-^ organoids showed an increase in immature cell types at the expense of most mature ExNs, as revealed by the percentage of cells per cluster (**Fig.S5G**) and visualization of cell density along pseudotime (**Fig.2D**, **Fig.S5H**). Thus, *ARID1B* mutations did not hinder dorsal telencephalic differentiation but altered the maturation trajectory of excitatory neurons at the transcriptional level.

**Figure 2.**
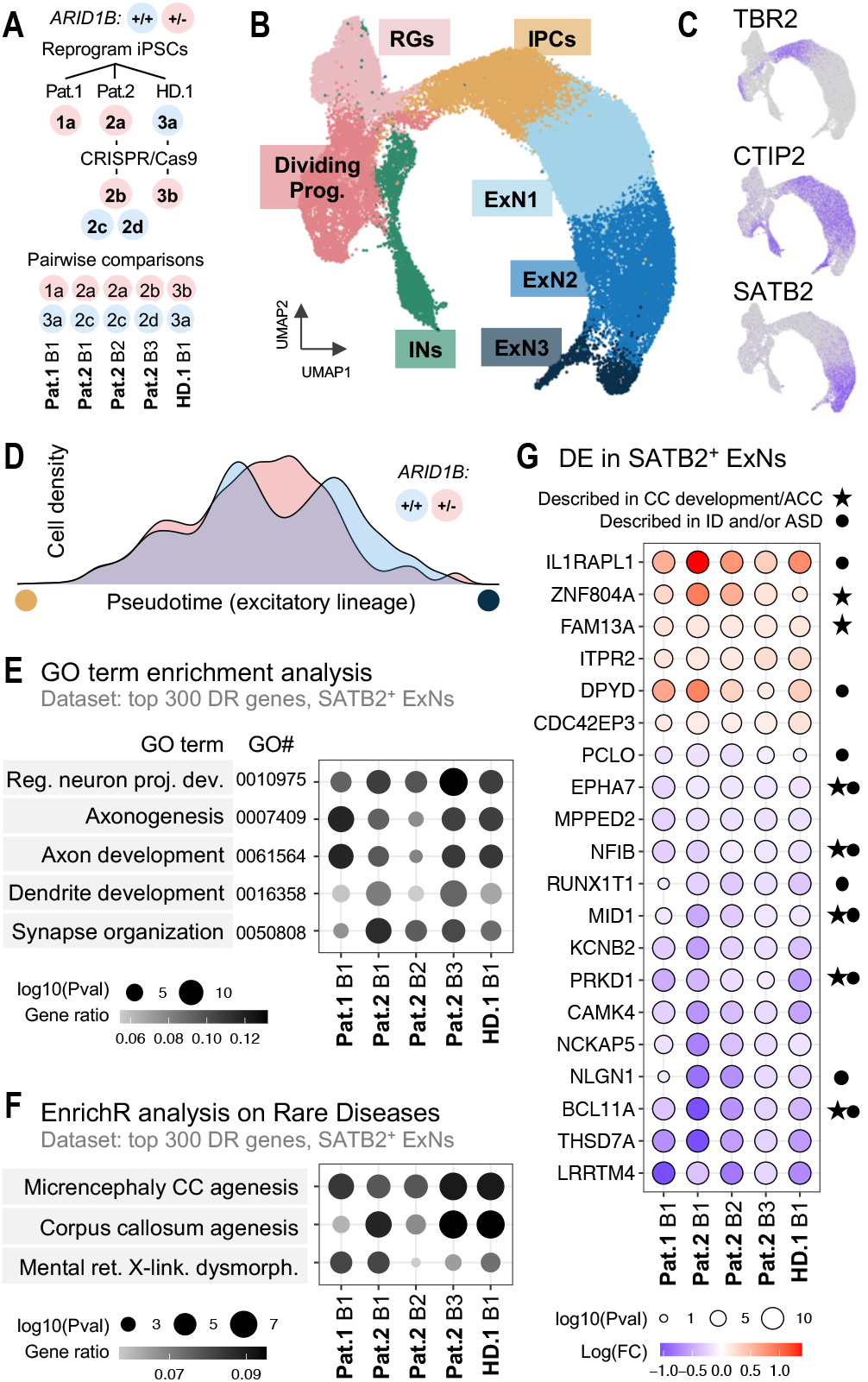
SATB2^+^ neurons in *ARID1B*^+/-^ organoids show dysregulation of genes associated with projection development and CC agenesis. **(A)** Cell line clones used for organoid generation and subsequent scRNAseq analysis at D120. Five independent pairwise comparisons were drawn between individually multiplexed organoids of different cell lines and batches; for Pat.2, two clones of *ARID1B*^+/+^ (2c, 2d) and *ARID1B*^+/-^ (2a, 2b) genotypes and three organoid batches (B1, B2, B3) were used. Comparisons were performed between organoids of the same batch, processed and sequenced simultaneously. **(B)** UMAP projection and unbiased clustering of 54.480 cells shows seven clusters of telencephalic populations: interneurons (INs), dividing progenitors, radial glia (RGs), intermediate progenitor cells (IPCs), and three populations of increasingly mature deep- and upper-layer excitatory neurons (ExN1, ExN2, ExN3) **(C)** TBR2, CTIP2 and SATB2 gene expression, marking different populations of cells from the excitatory lineage. **(D)** Slingshot analysis for pseudotime visualization of cell density along the excitatory lineage (from •IPCs to •ExN3), per genotype. **(E-G)** Analysis of differentially expressed (DE) genes in SATB2^+^ ExNs. Gene ontology (GO) term enrichment analysis of cell processes **(E)** and rare diseases **(F)** using the top 300 downregulated (DR) genes in *ARID1B*^+/-^ organoids, per pairwise comparison. The most statistically significant GOs are shown: GOs found in 5 out of 5 comparisons, with a gene ratio above 5%, and ordered by the average P value (Pval) across comparisons. **(G)** Up- and downregulated genes in *ARID1B*^+/-^ SATB2^+^ ExNs found in 5 out of 5 pairwise comparisons, ordered by descending fold-change (FC) (red, upregulated; violet, downregulated). Symbols mark previous associations with CC development or agenesis (ACC) (★) and intellectual disability (ID) or autism spectrum disorder (ASD) (•). Supporting literature: ID and/or ASD phenotypes have been associated mutations or encompassing microdeletions in *IL1RAPL1* (*46–48*), *DPYD* (*49, 50*), *PCLO* (*51*), *RUNX1T1* (*52, 53*), and *NLGN1* (*54*). Gene variants of *ZNF804A* are a risk factor for reduction in white matter in the CC (*55*); and *FAM13A* is a determinant of human brain asymmetry and handedness (*56, 57*). Callosal agenesis or dysgenesis has been described in patients harboring mutations or microdeletions in *EPHA7* (*58*), *NFIB* (*59, 60*), *MID1* (Opitz syndrome) (*61–64*), *PRKD1* (*65–67*), and *BCL11A/CTIP1* (2p15-16.1 syndrome) (*68–73*).

### Transcriptional programs of CC development are dysregulated in *ARID1B*^+/-^ SATB2^+^ neurons

To address specific effects on putative callosal projection neurons, we focused the subsequent analyses on cells in the ExN clusters that expressed SATB2 (SATB2^+^ ExNs, **Fig.2E- G**, **Fig.S6**). To group dysregulated genes by classes, we performed gene ontology (GO) term enrichment analyses on the top 300 genes downregulated in *ARID1B*^+/-^ organoids. The identified ontologies were linked to groups of genes with various overlaps across pairwise comparisons (**Fig.S6A-B**). Firstly, these genes were associated with cellular processes of projection neuron development, axonogenesis, dendrite development, and synapse organization (**Fig.2E**, **Fig.S6A**). Secondly, to explore the disease relevance of downregulated genes, we assessed the enrichment of terms from the Rare Disease AutoRIF database of Enrichr, which gathers almost 5 million gene-publication associations (*43–45*). Remarkably, disease ontologies involving corpus callosum agenesis were the most consistent and statistically significant across all five comparisons, followed by intellectual disability (**Fig.2F**, **Fig.S6B**). Together, these data suggest that the gene expression landscape of SATB2^+^ neurons is affected by *ARID1B* mutations, showing downregulation of broad transcriptional programs associated with neurite development and corpus callosum agenesis.

### Top-ranked genes dysregulated in *ARID1B*^+/-^ SATB2^+^ neurons control neuron projection development

To gain further insight into the specific genes dysregulated upon *ARID1B* haploinsufficiency, we focused on the most promising candidates – single genes consistently found upregulated or downregulated in SATB2^+^ neurons in 4/5 (**Fig.S6C-D**) or 5/5 pairwise comparisons (**Fig.2G**, **Fig.S6E-F**). We identified several ID and/or ASD risk genes (**Fig.2G**,•), as well as known players in CC development in humans (**Fig.2G**, ★). Interestingly, in addition to ID, CC abnormalities have been described in patients harboring mutations or microdeletions in *EPHA7* (*58*), *MID1* (Opitz syndrome) (*61–64*), *PRKD1* (*65–67*), *NFIB* (*59, 60*), *BCL11A/CTIP1* (2p15-16.1 syndrome) (*68–73*) and *AKT3* (*74*). Furthermore, some of the top-ranked dysregulated candidates have been shown to cause cell autonomous phenotypes in mouse neurons, such as reduced dendrite and axon formation in mouse hippocampal neurons upon IL1RAPL1 overexpression (*75*); defects in pioneering callosal projection neurons in *Nfib*-KO mice (*60, 76*); dysregulation of callosal axon outgrowth and mis-targeting in *Mid1*-KO mice (*77*); and strong defects on the morphology, migration, specification, or projection pattern of CPN subpopulations in *Bcl11a*-KO mice (*78–80*). Together, dysregulation of these genes may have synergistic effects on the development and projection of SATB2^+^ neurons in ARID1B patients. Given the overlapping effects between ACC and ID, we sought to precisely evaluate its impact on the formation of long-range projections.

### Late-stage telencephalic organoids form axon tracts *in vitro*

To assess if the identified transcriptional differences led to defects in the formation of long-range projections, we used an axon tract formation assay previously shown to recapitulate tract formation from early-stage organoids (*81*). Briefly, organoid pieces were dissected and placed in microdevices containing eight sets of two wells interconnected by a 7 mm-long Matrigel-coated channel (**Fig.3A**, **Fig.S7**). The assay started at D120 of organoid development and lasted 30-40 days. To visualize axon tract formation over time, we performed weekly brightfield imaging. Axon outgrowth initiated 24 h after the start of the assay, axon bundling after 4 to 5 days, and axons ultimately reached the contralateral side, forming a thick 7 mm-long tract after a total of 30 days (D120+30; **Fig.3B**, **Fig.S8A**). To assess the identity of projecting axons, we co-stained the pan-axonal marker Neurofilament (SMI312) (*82*), with markers known to be expressed in the corpus callosum *in vivo*, including Neuropilin-1 (NRP1) (*83–87*), Neogenin (NGN) (*88*), and Nectin-3 (*40, 89*). All markers were expressed at early (D120+7, **Fig.S8B**) and late (D120+30, **Fig.3C**, **Fig.S8C**) stages of tract formation across cell lines, supporting comparable identity of axonal projections. Thus, late-stage telencephalic organoids can be used to model long-range axon tract formation *in vitro*.

**Figure 3.**
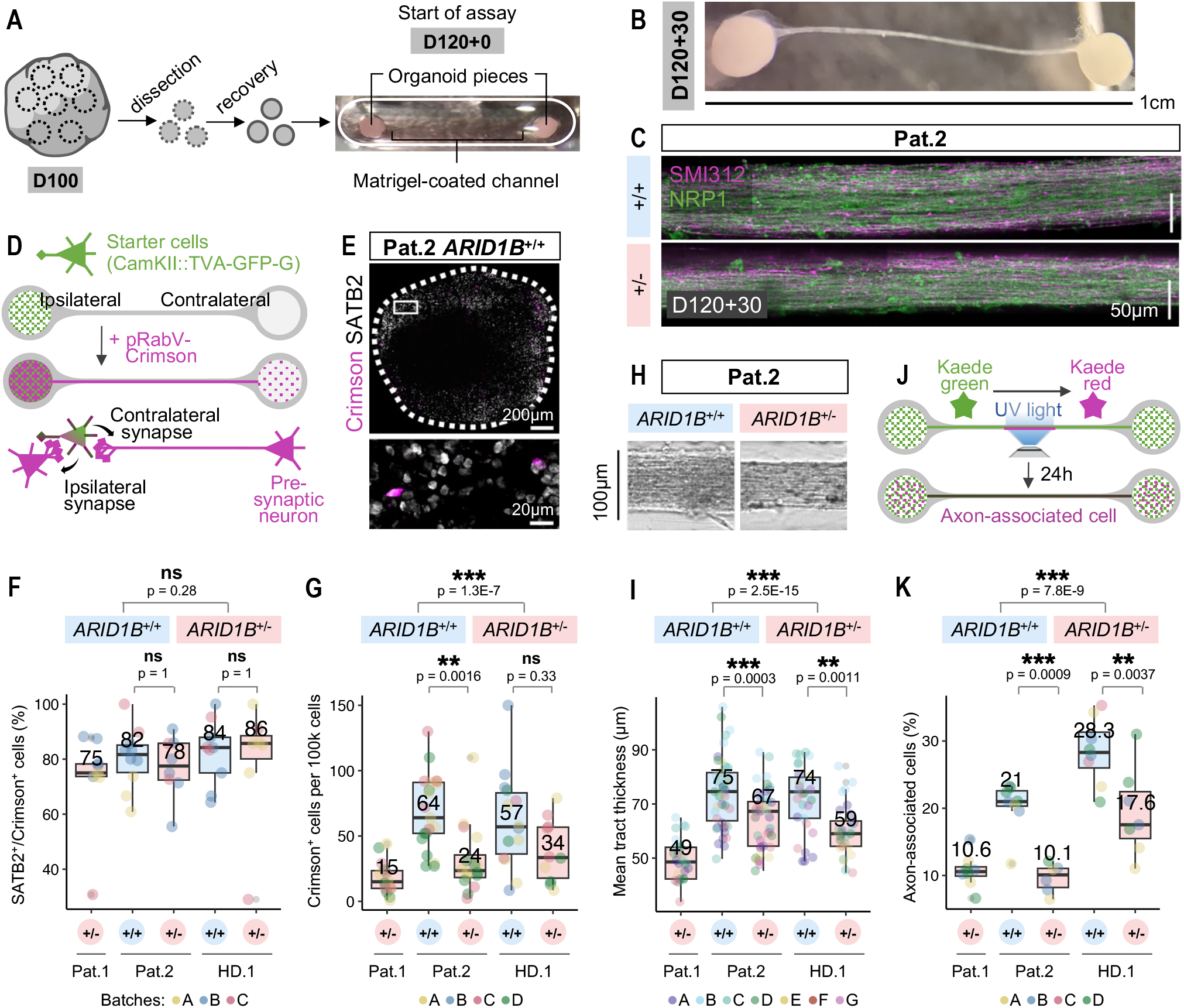
CC-like tracts from *ARID1B*^+/-^ organoids recapitulate underconnectivity phenotypes. **(A)** Schematic representation of the axon tract formation assay. Dissected organoid pieces are placed in two wells interconnected by a Matrigel-coated channel at D120 of development. **(B)** Imaging at 30 days after the start of the assay (D120+30); organoids are interconnected by an axon tract. **(C)** Immunostaining of axonal markers SMI312 and NRP1 at D120+30. **(D)** Schematic representation of monosynaptic pseudotyped rabies virus (pRabV) tracing, used to assess the formation of contralateral synapses. **(E)** Co-immunostaining of SATB2 and Crimson in a contralateral organoid (D120+30). **(F)** Quantification of the percentage of contralaterally connected cells (Crimson^+^) that are SATB2^+^, based on co-expression in immunostaining (D120+30). **(G)** Quantification of the absolute number of Crimson^+^ cells per 100k cells in contralateral organoids, identified by flow cytometry (D120+30). **(H)** Brightfield imaging of a segment of the final tract in Pat.2 organoids (D120+30). **(I)** Quantification of tract thickness (D120+30). **(J)** Schematic representation of Kaede photoconversion assay, used to assess the proportion of axon-associated cells. **(K)** Quantification of the percentage of axon-associated cells, identified by flow cytometry (D120+40). In boxplots, median values are indicated; each datapoint is an individual connected pair/organoid (technical replicate); datapoint colors indicate organoid batches (biological replicates). Statistical tests are analysis of variance (ANOVA); 0 ≤ p < 0.001, ***; 0.001 ≤ p < 0.01, **; 0.01 ≤ p < 0.05, *; p ≥ 0.05, ns (see results of statistical tests in **Table S4**)

### *ARID1B*^+/-^ CC-like tracts show reduced contralateral connectivity of SATB2^+^ neurons

To characterize the number and identity of neurons forming long-range connections, we used monosynaptic pseudotyped Rabies-virus (pRabV) tracing (**Fig.3D**, **Fig.S9A**). PRabV can enter starter cells (SCs) that express the cell-surface receptor TVA (here fused to GFP) and the rabies virus glycoprotein (G). Upon entry, pRabV can spread once retrogradely across structural synapses, to immediate pre-synaptic partners. For tract formation, only one of the two organoid pieces contained starter excitatory neurons, ensuring unilateral pRabV entry. Therefore, upon exposure to Crimson-expressing pRabV, Crimson^+^ cells in contralateral organoids were pre-synaptic neurons that connected through a long-range projection with an excitatory neuron. To assess the putative callosal identity of connected neurons, we co-stained Crimson and SATB2 (**Fig.3E**, **Fig.S9B**). Around 80% of contralateral connections were from SATB2^+^ neurons in *ARID1B*^+/+^ and *ARID1B*^+/-^ organoids (**Fig.3F**), showing an enrichment for callosal-like projections independent of genotype. Next, we interrogated the abundance of inter-organoid connections. Remarkably, flow cytometry analysis of contralateral Crimson^+^ cells showed a significant reduction in the absolute number of contralateral structural synapses in *ARID1B*^+/-^ connected organoids (**Fig.3G**). Thus, *ARID1B* mutations caused a decrease in contralateral connectivity after long-range projection of a population of neurons enriched in SATB2^+^ CPNs.

### CC-like tracts from *ARID1B*^+/-^ organoids have fewer long-range projections

To evaluate if decreased connectivity was due to fewer axons or lower percentage of cells forming synapses after projection, we assessed axon tract properties. Axon outgrowth and bundling occurred in *ARID1B*^+/+^ and *ARID1B*^+/-^ organoids (**Fig.3C**, **Fig.S8C**), ruling out obvious defects in axon fasciculation, which are responsible for CC defects in other disease settings (*90*). However, the thickness of the interconnecting axon tract was significantly reduced in *ARID1B*^+/-^ organoids (**Fig.3H-I**, **Fig.S8D**). To probe for differences in tract compaction, we assessed the proportion of neurons contributing axons to the tract. We resorted to mosaic expression of the photoconvertible protein Kaede, whose fluorescence can be converted from green to red by exposure to UV light (**Fig.3J**, **Fig.S9C**). Photoconversion restricted to the central tract region (**Fig.S9D**) led to local formation of Kaede-red, which diffused to the associated cell bodies within 24 h. Analysis by flow cytometry one day after photoconversion showed a significantly lower percentage of axon-associated cells (Kaede-red^+^) in *ARID1B*^+/-^ organoids (**Fig.3K**, **Fig.S9E-F**). Together, these observations indicate a reduction in the number of axons interconnecting *ARID1B*^+/-^ organoids, compared to control.

### Axonogenesis is impaired in SATB2^+^ projection neurons of *ARID1B*^+/-^ organoids

We hypothesized that reduced tract thickness in *ARID1B*^+/-^ organoids could be either due to slower axonogenesis or to a lower number of neurons forming long axonal projections. To test these hypotheses, we analyzed early stages of tract formation, prior to the onset of axon fasciculation and bundling (**Fig.4A**). Axon outgrowth properties were extracted from live imaging performed between 24 h and 72 h after the start of the assay (**Fig.4B**). To quantify axon outgrowth, we developed an automated analysis pipeline for axon segmentation and counting (**Fig.S10A**, details in **Materials and Methods**). Notably, *ARID1B* mutations caused a significant reduction in the number of axons observed at 72h (**Fig.4C**, **Fig.S10**), as measured by quantification of axon number near the organoid (first 160 µm, **Fig.4D**), and along the initial segment of the channel (**Fig.4E**). To assess whether this reduction could be a result of lower outgrowth speed, we used manual tracking of individual growth cones. Axon outgrowth speed averaged at 37µm/h and was unchanged between *ARID1B* genotypes of isogenic backgrounds, showing only a slight cell-line-dependency (**Fig.4F**). These results indicate that *ARID1B* haploinsufficiency did not affect the axon outgrowth machinery itself but caused a significant reduction in the number of long-range projections at early stages of CC-like tract formation.

**Figure 4.**
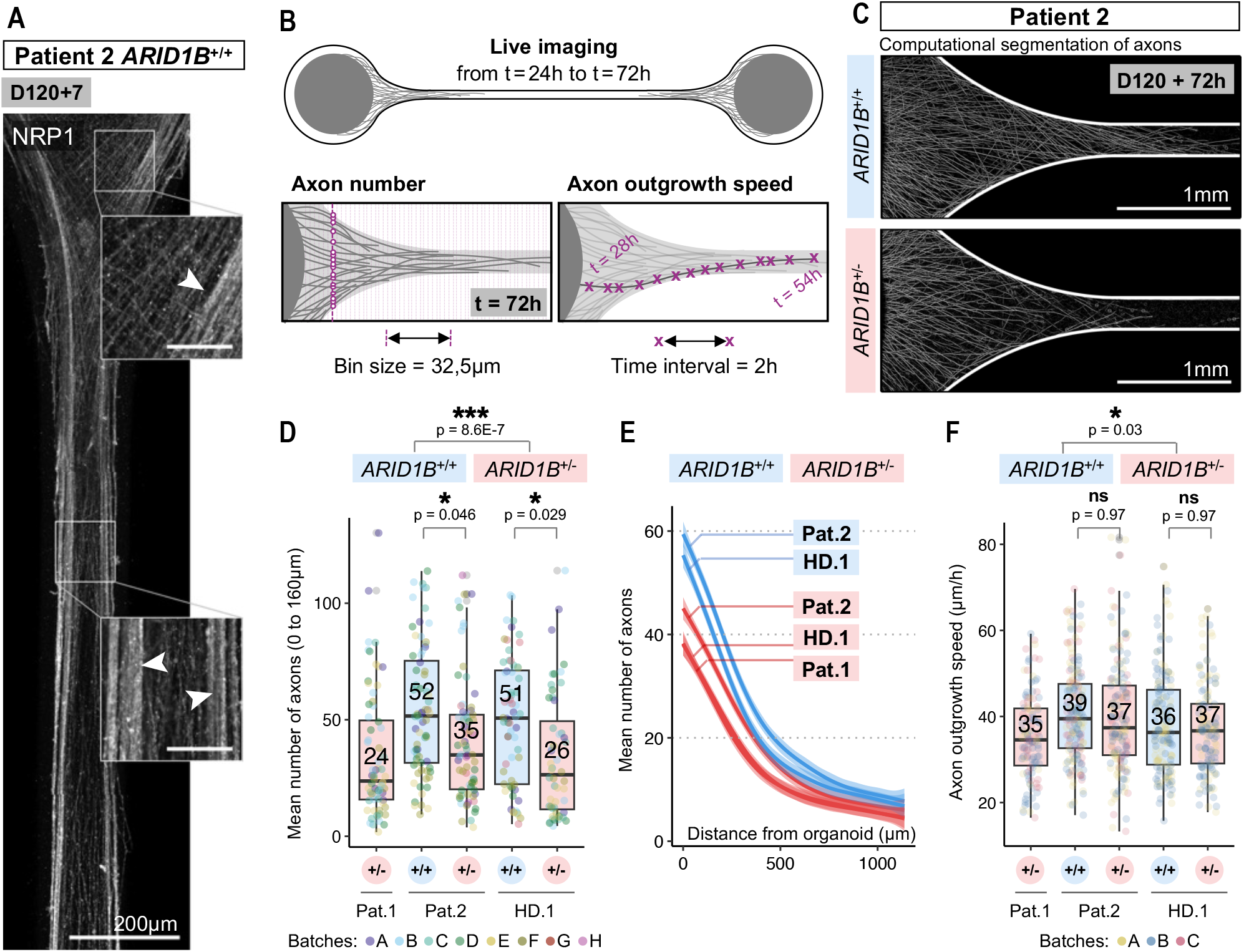
A lower proportion of neurons send long-range axonal projections in *ARID1B*^+/-^ organoids. **(A)** Immunostaining of the CC marker NRP1 at D120+7. Note the start of axon bundling at the edges of the channel (arrowheads, ➤) **(B)** Live imaging of early stages of axon outgrowth allows the quantification of number and speed of growing axons. Axon number was measured at every 32,5 µm (50 pixels); and outgrowth speed was measured by manual tracking of individual growth cones with an imaging frequency of 2 hours. **(C)** Representative images of computational segmentation of axons at D120+72h. **(D-E)** Quantification, at D120+72h, of the mean number of axons in close proximity to the organoid (each datapoint is an individual organoid) **(D)**; and across the initial segment of the channel **(E)**. **(F)** Quantification of axon outgrowth speed (each datapoint is an individual axon). In boxplots, median values are indicated; each datapoint is an individual organoid/axon (technical replicate); datapoint colors indicate organoid batches (biological replicates). Statistical tests are analysis of variance (ANOVA); 0 ≤ p < 0.001, ***; 0.001 ≤ p < 0.01, **; 0.01 ≤ p < 0.05, *; p ≥ 0.05, ns (see results of statistical tests in **Table S4**).

### Long-range projections of subcerebral projection neurons are unaffected in *ARID1B*^+/-^ organoids

Finally, to assess the specificity of the identified underconnectivity phenotypes to callosal projections, we evaluated tract formation at early stages of organoid development (D30 to D60) (**Fig.S11**). At these stages, SATB2^+^ neurons were virtually absent (**Fig.S11A**) and organoids were mostly composed of early-born deep-layer TBR1^+^ and CTIP2^+^ neurons (**Fig.S3A**, **Fig.S11A**), which project to subcerebral targets *in vivo* (*91–93*). To evaluate early axon outgrowth, we performed live imaging. Axon outgrowth speed averaged at 28µm/h, independently of *ARID1B* genotype (**Fig.S11B**). Thus, *in vitro*, subcerebral classes of projection neurons extended axons at a lower speed than callosal projection neurons (**Fig.4F**), which may reflect properties related to differential axon outgrowth characteristics *in vivo* (*94, 95*). Next, to evaluate tract formation, we performed imaging at D30+30 (**Fig.S11C**). Remarkably, the thickness of *ARID1B*^+/-^ tracts was not reduced compared to isogenic controls at this stage (**Fig.S11D-E**). Thus, *ARID1B* haploinsufficiency did not affect the proportion of subcerebral projection neurons capable of sending long-range projections *in vitro*. This result is in stark contrast to the effects described for SATB2^+^ projection neurons (D120+30, **Fig.3H-I**), supporting the *in vitro* recapitulation of specific white matter phenotypes of ARID1B patients.

## Discussion

Heterozygous mutations in chromatin remodelers, including mSWI/SNF complex members, cause severe neurodevelopmental defects in humans (*4*). However, due to the inaccessibility of human brain tissue, the entangled neurodevelopmental phenotypes seen in patients, and difficulties in cross-species modelling of brain development, the cellular and molecular origin of these conditions have remained poorly understood and challenging to study.

Here, we conducted a holistic and quantitative comparison of several processes of CPN biology between *ARID1B*^+/+^ and *ARID1B*^+/-^ human neural organoids. We show that *ARID1B* haploinsufficiency does not affect fate-acquisition of SATB2^+^ neurons. However, it causes transcriptional dysregulation of genes involved in neuron projection development, leading to a significant reduction in the formation of long-range callosal projections and resulting in broad structural underconnectivity phenotypes. Transcriptome data further suggest that *ARID1B* mutations may impact CC development by additional mechanisms, such as impaired axon guidance due to *EPHA7* downregulation (*96–98*) or midline glia abnormalities due to *NFIB* downregulation (*60, 76, 99, 100*). Therefore, our study expands on known functions of the mSWI/SNF complex in the control of neuronal morphology, neurite formation, and axon fasciculation and pathfinding (*101*–*106*). We uncover a role of ARID1B in the regulation of global transcriptional programs required for human corpus callosum development, and specifically needed for axon projection formation in callosal projection neurons.

Thus far, mouse models have been the gold standard to understand ACC. However, while CC development spans around 30 weeks of gestation in the human, it lasts just a few days in the mouse (*15, 89, 107, 108*). Such inter-species difference may explain why phenotypes observed in heterozygosity in humans often require homozygous loss of functional alleles in mice, as seen in several mSWI/SNF members (*109, 110*). Also, in the presence of severe midline abnormalities *in vivo* (*111, 112*), it becomes challenging to identify additional cell-autonomous impairments at the level of callosal projection neurons. Here, we show that neural organoids are superior to mouse models in mimicking the protracted timeline and genetic context of human development, not only recapitulating the maturation trajectory of CPNs, but also revealing disease features in heterozygosity. Furthermore, we develop a robust and reproducible method to probe differentiation, maturation, axonogenesis, axon fasciculation, and synapse formation processes in human CPNs, thereby isolating phenotypes that could be overlooked in a complex *in vivo* environment. Finally, whereas *Arid1b*^+/-^ mice do not replicate the complete ACC observed in patients, they exhibit a significant reduction in cortex size (*28*), a feature that is not recapitulated in the human cortex. These discrepancies suggest species-specific functions of ARID1B in cortical and corpus callosum development that could only have been identified with human-specific *in vivo* and *in vitro* studies. Thus, a re-examination of ACC phenotypes in the context of human neurodevelopment may provide new insights that were previously inaccessible, contributing to a more complete understanding of these conditions.

On par with corpus callosum agenesis, mutations in mSWI/SNF complex members are strongly associated with syndromic forms of intellectual disability and autism spectrum disorder (*4–6, 113–115*). The question of whether and how ACC and ID/ASD phenotypes are related remains central to this field. The prevailing hypothesis, based on mouse models, is that a decrease in interneuron production and inhibitory synapses causes ID/ASD (*29, 30, 116*). However, recent neural organoid studies, including our own, challenge the generalization of these mechanisms to human development, by demonstrating an expansion of ventral telencephalon lineages upon *ARID1B* haploinsufficiency (*32, 33*). Furthermore, the evolution of human cognition highly correlates with cortical expansion and increased number, diversity, and complexity of cell types in the upper cortical layers (*89, 108, 117–121*). Accordingly, while increased synaptic processing capacity of upper-layer neurons has been associated with higher intelligence (*119*), excitatory synapse pathophysiology results in ASD and ID (*122*). In this context, we show that multiple genes found dysregulated in SATB2^+^ neurons are associated with ID and ASD. Also, several differentially expressed genes participate in excitatory synapse formation, such as *KCNB2* (*123*), *CAMK4* (*124*), *LRRTM4* (*125, 126*), *NLGN1* (*127*), *PCLO* (*51, 128*), and *IL1RAPL1* (*129–132*). Thus, excitatory/inhibitory imbalance, as well as transcriptional dysregulation in upper-layer neurons, may impact higher-order processing functions and contribute to the cognitive and behavioral phenotypes of ARID1B patients. Overall, our data suggest that coexisting ID/ASD and ACC are under the control of distinct transcriptional and cell biological mechanisms, shedding new light onto the multifactorial nature of the ARID1B phenotype.

Corpus callosum agenesis is one of the most prevalent clinical findings in prenatal MRIs, with highly unforeseeable prognosis (*133–135*). Although previous studies have shown favorable outcomes in approximately two thirds of patients (*135, 136*), prenatal diagnosis of ACC is a surging cause of pregnancy termination (*133*). Pinpointing affected cell populations and biological processes responsible for this condition, developing appropriate models that mimic disease features while preserving a human genetic context, and distinguishing structural from cognitive phenotypes have been long-standing goals with transversal medical and biological relevance. After decades of clinical studies showing a correlation between CC malformations and *ARID1B* mutations (*3, 6, 25*), our study provides an unprecedented leap in the molecular and cellular understanding of this pathology and unlocks a paradigm shift in human corpus callosum development research.

## Acknowledgments

We thank all members of the Knoblich lab for fruitful discussions and feedback on the manuscript. We thank Dr. Tatsuya Osaki for providing the Kaede photoconversion protocol. We thank the IMBA stem cell core facility for cell reprogramming services; the IMBA/IMP/GMI BioOptics facility for microscopy services; the Next Generation Sequencing Facility at Vienna BioCenter Core Facilities (VBCF) for single cell sequencing services; the IMBA/IMP/GMI Bioinformatics for sequencing analysis; the IMBA/IMP/GMI Histopathology facility for cryosectioning and immunostaining of fetal brain samples; and A. Meixner for coordinating ethical approvals. We especially thank all patients and their families for participating in this study or donating tissue.

## Funding

Work in the Knoblich laboratory is supported by the Austrian Academy of Sciences, the Austrian Science Fund (FWF), (Special Research Programme F7804- B and Stand-Alone grants P 35680 and P 35369), the Austrian Federal Ministry of Education, Science and Research, the City of Vienna, and a European Research Council (ERC) Advanced Grant under the European Union’s Horizon 2020 programs (no. 695642 and no. 874769). JS was supported by the EMBO long term fellowship (EMBO ALTF 794- 2018) and the European Union’s Horizon 2020 research and innovation program under the Marie Skłodowska-Curie fellowship agreement 841940. Work in the Ikeuchi laboratory is supported by JSPS (20K20643 and 20H05786), AMED (JP20gm1410001), and Institute for AI and Beyond.

## Author contributions

Conceptualization: CMC, JS, GK, DP, YI, NSC, JAK; Methodology: CMC, VP, AW, BD, TL, MP, AP, PM, MS, SYAC, RS, RH, GK, YI; Investigation: CMC, AW, MN, MS, GK; Visualization: CMC; Funding acquisition: JAK, YI; Project administration: JAK; Supervision: JS, NSC, JAK; Writing – original draft: CMC; Writing – review & editing: CMC, NSC, JAK, with input from all authors.

## Competing interests

JAK is inventor on a patent describing cerebral organoid technology and co-founder and scientific advisory board member of a:head bio AG.

## Data and materials availability

The scRNAseq data discussed in this publication have been deposited in the European genome-phenome Archive (EGA) and in the NCBI’s Gene Expression Omnibus (*137*) and are accessible through GEO Series accession number GSE231546. Analysis was performed as outlined in the **Materials and Methods**. No custom code was generated for this study; used code is available upon request. ARID1B patient iPSCs will be made available upon request after obtaining ethical approval from the Ethics Committee of the medical university of Vienna (MUV) under a materials transfer agreement with the Institute of Molecular Biotechnology of the Austrian Academy of Sciences. This study was approved by the local ethics committee of the MUV.

## Materials and Methods

### Patient sample selection

#### Inclusion criteria

The study was approved by the local ethics committee of the Medical University of Vienna (MUV). Study inclusion criteria were as follows: 1) complete or partial corpus callosum agenesis proven by fetal MRI; 2) *ARID1B* mutations proven by genetic testing with whole exome sequencing; 3) continuous follow-up at the department of Pediatrics at the MUV.

#### Tissue sample collection

All clinical data were derived from the MUV patient registry (e.g. sex, age, pre- and postnatal MRIs and DTIs). After informed consent, 10 mL blood was collected from two selected patients for iPSC reprogramming. Evaluation of formalin-fixed and paraffin embedded (FFPE) brain material of one other patient and an age-matched control case was performed. Fetal MRI of several patients and control cases was included for comparative analysis.

### Fetal MRIs

#### Inclusion criteria

Women with singleton pregnancies undergoing fetal MRI at a tertiary care center from January 2016 to January 2023 were retrospectively reviewed. A retrospective IRB-approved review of patient records undergoing clinically indicated fetal MRI was performed and patients with confirmatory genetic testing report for *ARID1B* mutation were selected. Gestational ages at the time of fetal MRI (given in gestational weeks and days post menstruation) were determined by first-trimester ultrasound. High-quality super-resolution reconstruction was obtained. Age-matched control cases without *ARID1B* mutations were identified and selected based on an absence of confounding comorbidities, including structural cerebral or cardiac anomalies and fetal growth restriction.

#### Measurement of cortical volume

Fetal MRI scans were conducted using 1.5 T (Philips Ingenia/Intera, Best, Netherlands) and 3 T magnets (Philips Achieva, Best, Netherlands). The examinations were performed within 45 min and both the fetal head and body were imaged. Fetal brain imaging included T2-weighted sequences in three orthogonal planes (slice thickness 3-4 mm, echo time = 140 ms, field of view = 230 mm) of the fetal head. Post-processing was conducted in a similar methodology as previously described (*134, 138, 139*). Super-resolution imaging was generated using a volumetric super-resolution algorithm (*140*). The resulting super-resolution data were quality assessed and only cases that met high-quality standards (score ≤ 2 out of 5) were included in the analysis. Atlas-based segmentation (*138*) of the cortex was performed using the open-source application ITK-SNAP (*141*). Volumetric data was calculated based on the investigated gestational ages. Plotting was done in R with the geom_smooth() function of ggplot2, using method “lm” and formula y ∼ x + I(x^2^).

### Generation of iPS cells

Induced pluripotent stem cells were generated from peripheral blood mononucleated cells (PBMCs) isolated from patient blood samples as previously described (*142*). Briefly, 10 mL blood was collected in sodium citrate collection tubes. PBMCs were isolated via a Ficoll-Paque density gradient and erythroblasts were expanded for 9 days. Erythroblast-enriched populations were infected with Sendai Vectors expressing human OCT3/4, SOX2, KLF4, and cMYC (CytoTune, Life Technologies, A1377801). Three days post-infection, the cells were switched to mouse embryonic fibroblast feeder layers; 5 days post-infection, the medium was changed to iPSC media (KOSR + FGF2); and 10 to 21 days post-infection, the transduced cells began to form colonies that exhibited iPSC morphology. IPSC colonies were picked and passaged every 5 to 7 days. IPSCs were passaged 10 times before being transferred to the mTeSR1 culture system (StemCell Technologies, 85875).

### *ARID1B* mutations identified by genome sequencing

A schematic representation of the ARID1B protein domains and location of mutations is depicted in **Fig. 1C**.

Patient 1 presented a microdeletion (6q25.3del) encompassing part of *ARID1B* (from exon 8) and two downstream genes (*TMEM2432* and partially *ZDHHC14*), identified clinically by whole exome sequencing and confirmed in iPSCs with single nucleotide polymorphism (SNP) genotyping.

Patient 2 harbored a heterozygous single-nucleotide duplication (NM_020732.3: c.2201dupG) that led to an early stop codon in exon 8, upstream of the active domains of the ARID1B protein, identified clinically by whole exome sequencing and confirmed in iPSCs with Sanger sequencing.

The engineered cell line, HDon.1 *ARID1B*^+/-^, harbored a heterozygous mutation in the ARID domain of *ARID1B* (NM_020732.3: c.3489_3490insNN), which led to the formation of an early stop codon in exon 14, identified with Sanger sequencing.

### Genome engineering of iPSCs

#### Generation of isogenic control cell lines for Patient 2

An isogenic control cell line of Patient 2 was generated using CRISPR/Cas9. S. pyogenes Cas9 protein with two nuclear localization signals was purified as previously described (*143*). The transcription of guide RNAs (gRNAs) was performed with HiScribe T7 High Yield RNA Synthesis Kit (NEB) according to the manufacturer’s protocol, and gRNAs were purified via Phenol:Chloroform:Isoamyl alcohol (25:24:1; Applichem) extraction followed by ethanol precipitation. The sequence of the sgRNA used was 5’ GGGGGGgCCCATCTCCCTC 3’ (g is the single-nucleotide duplication in Pat.2). The HDR template (custom ssODN, Integrated DNA Technologies) was designed to span 100 bp up and downstream of the mutation site. For generation of isogenic control cell lines, iPSCs were grown in mTeSR1. Cells were washed with PBS (pH 7.4, without MgCl_2_ or CaCl_2_; Gibco, 14190-250) and incubated in TrypLE Select (Thermo Fisher Scientific) for 5 min at 37 °C. After formation of a single cell suspension, cells were resuspended in mTeSR and viability was assessed. Cells were washed in PBS and 1 x 10^6^ cells were resuspended in Buffer R of the Neon Transfection System (Thermo Fisher Scientific). 12 ng of sgRNA and 5 ng of electroporation- ready Cas9 protein were mixed to form the Cas9/sgRNA RNP complex and the reaction was mixed and incubated at 37 °C for 5 min. 1 µL of the HDR template (500 µM) was added to the Cas9/sgRNA RNP complex and combined with the cell suspension. Electroporation was performed using a Neon® Transfection System (Thermo Fisher Scientific) with 100 µL Neon® Pipette Tips using the ES cells electroporation protocol (1400 V, 10 ms, 3 pulses). Cells were seeded in one well of a 6 well plate in mTeSR1 supplemented with Rho-associated protein kinase (ROCK) inhibitor (1:100; Selleck Chemicals, S1049). After a recovery period of 5 days, cells were plated at a low density in a 10 cm dish, to form separate colonies. Single colonies were picked after 5-7 days and transferred to 96 well-plates. After expansion, gDNA was extracted using the DNA QuickExtract Solution (Lucigen), followed by PCR and Sanger sequencing to determine efficient mutation rescue.

#### Generation of *ARID1B* mutation in a healthy donor background

For generation of an *ARID1B* mutation in a cell line derived from a healthy donor (HD.1 *ARID1B*^+/-^), we targeted the ARID domain with CRISPR/Cas9. The procedure was as described for the generation of the isogenic control cell line for Patient 2, with the exception that no HDR template was provided. The sequence of the gRNA used was 5’ CTGGCAACCAACCTAAACGT 3’. A heterozygous mutation on top of the ARID domain was generated by non-homologous end joining.

### Assessment of pluripotency and genomic integrity of iPSCs

#### Pluripotency

Pluripotency of cell lines was assessed by flow cytometry for stem cell markers SSEA-4 and TRA-1-60. Briefly, a single cell suspension was washed with PBS and fixed with 4% paraformaldehyde (PFA; Sigma-Aldrich, 441244-1KG) and 0.1% Saponin (Sigma-Aldrich, 47036- 50G-F), for 15 min at RT. Cells were stained with Alexa Fluor® 647 Mouse anti-SSEA-4 (Clone MC813-70) (BD Biosciences, 560796) and Alexa Fluor® 488 Mouse anti-Human TRA-1-60 (Clone TRA-1-60) (BD Biosciences, 560173) (1:20), in PBS with 0.1% Saponin and 1% BSA, for 30 min on ice. Unstained cells were used as a control. Cells were washed PBS with 0.1% Saponin and 0.2% BSA and analyzed on a BD LSRFortessa Instrument (BD Biosciences). Single, live cells were gated, and the percentage of positive cells was analyzed using the FlowJo Software.

#### Genomic integrity

Genomic integrity of cell lines was analyzed by SNP genotyping (*144*), on an Infinium PsychArray v1.3 (Illumina) and compared to data from PBMCs. Normal karyotype was confirmed resorting to short tandem repeat (STR) analysis.

After quality control, cells were banked at different passages. For experiments, cells were used from passage 35 to 60, and never more than 10-15 passages after defrosting of a quality-controlled vial. At the end of each round of experiments, *ARID1B* mutations and genomic integrity were once again confirmed.

### Maintenance and passaging of iPSCs

All cells were cultured on 6 well-plates (Corning, 3516) coated with hESCs-qualified Matrigel (Corning, 354277), and maintained in complete mTeSR1 medium . Cells were fed daily with 2 mL of mTeSR1 (feeding with 4 mL allowed skipping of one feeding day per week) and passaged after reaching 60-80% confluency (every 3-5 days). For passaging, cells were exposed to 0.5 mM EDTA diluted in PBS during 3 min at 37 °C, lifted in mTeSR1 by gentle spraying of the bottom of the well, and triturated to small clusters of 20-50 cells. Cells were routinely tested for mycoplasma. Cells, embryoid bodies (EBs), and organoids were kept in a 5 % CO_2_ incubator at 37 °C.

### Generation of dorsal tissue-enriched telencephalic organoids

Media formulations can be found in **Table S1**. Feeding volumes in 96WP format were of 150 µL, and in 10 cm dish format of 15 mL. The protocol was as previously described ((*38*); Matrigel-free conditions), with the only exception that EBs were generated from 12.000 cells instead of 9.000 cells.

#### Embryoid body formation

Human iPSCs were grown to 60-80% confluency and dissociated into a single cell suspension by Accutase (Sigma-Aldrich, A6964) treatment for 5 min at 37 °C, followed by manual trituration. Cells were seeded in an ultra-low binding 96-well plate (Szabo-Scandic, COR7007), at a density of 12.000 live cells/well, in 150 µL of complete Essential 8 (E8) medium (Thermo Fisher Scientific, A1517001) with 50 μM ROCK inhibitor. On day 3, the medium was replaced with E8 without ROCK inhibitor supplementation. From day 6, EBs were fed daily with Neural Induction (NI, **Table S1**) medium.

#### Tissue patterning

On day 13, the NI medium was replaced with Differentiation Medium without vitamin A (Diff-A, **Table S1**). Two pulse applications of 3 µM of GSK-3 Inhibitor CHIR99021 (Merck Millipore, 361571) were done on days 13 and 14; the medium was replaced with Diff-A without CHIR on day 16 and exchanged daily until day 20.

#### Long-term culture

On day 20, the organoids were transferred to 10 cm dishes and cultured on an orbital shaker (Celltron) at a rotating speed of 57 rpm. On day 25, the medium was replaced with Differentiation Medium with vitamin A (Diff+A, **Table S1**). From day 25 onwards, the medium formulation remained unaltered throughout organoid development, and the medium was changed twice a week (every 3-4 days).

#### Organoid patterning confirmation

Patterning of organoids was verified at D60 resorting to SOX2, TBR2 and CTIP2 staining and at D90 resorting to SATB2 immunostaining. Batches in which dorsal cortical neural rosettes were present at early stages, and SATB2^+^ neurons were abundant around the whole organoid surface at late stages were used for further analyses.

### Virus production

The following plasmids were used in this study:

1. pAAV-CaMKII-TVA/EGFPfusion-N2cG-WPRE3-SV40pA.
2. pAAV-CAG-Kaede.
3. rAAV2-Retro-Helper (Addgene #81070) – AAV Retro Capsid.
4. pAdDeltaF6 (Addgene #112867) – AAV Helper Capsid.
5. CVS-N2C-Crimson (Addgene #172377).
6. pTIT-N, P, L (gift from the Molecular Biology Service at the ISTA, Institute of Science and Technology Austria).

#### AAV production

Adeno-Associated Virus (AAV) was generated as described by previous protocols (*145, 146*). Briefly, 10x15cm dishes of HEK293-GT cells were transfected with plasmids carrying the capsid, helper, and cargo genes according to previous methods (*145*). AAVs from the supernatant and cells were recovered, concentrated and iodixanol purified as described before (*146*). The purified virus was aliquoted and stored at −70 °C. qPCR was used to titrate the virus, as described previously (*147*).

#### Kaede plasmid production

To generate the AAV-CAG- Kaede plasmid, cDNA of Kaede (*148*) was cloned from CoralHue® Kaede (pKaede-MN1), together with the CAG promoter, into a pAAV plasmid.

#### Pseudotyped monosynaptic Rabies virus (RabV) production

EnvA pseudotyped rabies virus (pRabV) was produced as previously published (*149*), with minor modifications. Briefly, HEK293-GT cells were transfected with CVS-N2C-Crimson viral genome, pTIT-N, pTIT-P and pTIT-L using Xfect Transfection Reagent (Takara, 631318) according to the instructions of the manufacturer. The medium was changed on transfected cells every day for 4 days. The medium was not changed on day 5 and on day 6 but was collected, filtered, and used in the pseudotyping step. To pseudotype the virus, the media harvested from day 5 and 6 was applied to BHK-eT cells for 6h. Cells were then processed according to a previous protocol (*149*), until the step of purification of the pseudotyped pRabV. To concentrate the produced virus, the media collected from the BHK-eT cells were pooled, filtered, and incubated with Benzonase Nuclease (Merck Millipore, US170664-3) for 30 min at 37 °C. The medium was then ultracentrifuged on a sucrose cushion for 2 h at 70,000 rcf and removed from the viral pellets. Pellets were resuspended and pooled in 200 µL of PBS. Functional titer was determined as described before (*150*). The virus was aliquoted and stored at −70 °C until use; single aliquots were defrosted a maximum of 2 times.

### Axon tract formation assay

#### Microdevice fabrication

The microdevices for axon tract formation assay were fabricated as previously described (*151*).

#### Organoid dissection

Organoids were dissected at D95±5, as depicted in **Fig.S7C**. P1000 tips were cut, generating an opening with a width of around 1.5 mm. Organoids were punctured with the cut tips, forming round pieces. Each organoid was used to generate 5 to 10 pieces; 4 to 5 organoids were dissected per batch and per cell line, to yield around 30 optimally sized pieces, with 1-1.5mm in diameter. For tract formation at early developmental stages, this step was not necessary, as D30 organoids were still small enough to fit into the microdevice wells.

#### Microdevice preparation

Microdevices were kept submerged in 96% ethanol until use. One day before the start of the assay, ethanol-covered microdevices were placed on top of slides and pressed down firmly. Nunc® Lab-Tek™ Chambered Coverglasses with 1 well (Thermo Fisher Scientific, 155361PK) were used for experiments involving live imaging of axon outgrowth or immunostaining and confocal imaging; Permaflex Plus Adhesive Microscope Slides (Leica, 3830001) were used for all other experiments. Microdevices were allowed to completely dry and attach to the slides overnight. Microdevice channels were coated with 5% Matrigel for 2 h at 37 °C, as previously described (*81, 151*). Briefly, the coating solution was sucked with a P1000 pipette from one well to the other, through the channel, filling it up completely and without air bubbles.

#### Assay setup

The assay started at D120 (±5 days) or at D30 (±4 days) of organoid development, as previously described (*81, 151*). Two organoids of the same genotype and same batch were placed in opposing wells, connected by a Matrigel-coated microchannel. Organoid pieces were chosen based on optimal size to fit into the wells (1-1.5mm in diameter). Placement of organoid pieces in the wells marked D120+0 (or D30+0) in the assay.

#### Feeding

Microdevices were fed on day1-2 and then twice a week (every 3 to 4 days). Feeding medium was Diff+A + 2% Matrigel on day 0-7, Diff+A + 1% Matrigel on day 7-14, and Diff+A without Matrigel thereafter.

#### Rabies virus tracing

For tracing of contralateral synapses, the protocol was as depicted in **Fig.S9A**. Organoid pieces were exposed to AAV::CamKII-TVA-GFP-G to generate starter cells. Three rounds of infection were performed, each with 0.125 µL of virus (MOI = 6,6E+13) per plate. Typically, 10 to 15 organoid pieces from the same batch were exposed together in a 6 cm dish with 4 mL of Diff+A medium. The first infection was done at D110 (±5 days); the second infection was done 2-3 days later (D112), without medium change (just virus addition); the third infection was done at D116 with medium change and virus addition; after 3-4 days, the medium was replaced with plain Diff+A and pieces were ready for the assay. These successive infections significantly increased the number of starter cells generated, in comparison with a single infection. The axon tract formation assay was setup with only one of the two organoids expressing TVA and G and, therefore, permissive for Rabies virus entry. As negative control, 1-2 tracts per device were formed with two wild-type pieces. Exposure to pRabV occurred at day D120+20 (±1 day) and organoids were analysed (by fixation and immunofluorescence, or by flow cytometry) 10 days later, at D120+30. At this stage, abundant ipsilateral and sparse contralateral Crimson^+^ cells were visible under a widefield cell culture microscope; no Crimson^+^ cells were seen in negative control samples, as expected.

#### Kaede tracing

For assessment of axon-associated cells, the protocol was as depicted in **Fig.S9C**. Organoids were exposed to AAV::CAG-Kaede-green on D120+21, by addition of 0,127 µL of virus (MOI = 6,0E+13) per well (200 µL of medium). Kaede photoconversion was done at D120+39. The wells were covered below with black tape, leaving only the central part of the axon tracts exposed. This helped avoid unwanted conversion of cell bodies. After, the microdevices were placed in an inverted Scanning confocal microscope LSM800 (Zeiss). Each axon tract was exposed to 15min of blue UV light, with a 10x objective, leading to the local formation of Kaede-red. As negative control, one tract per device was left unconverted. After conversion, the medium was exchanged and the microdevices were placed in the incubator for another 24 h, during which there was diffusion of Kaede-red to the cell bodies of axon-associated cells.

#### Fixation

Axon tracts were fixed inside the microdevices. After one wash with PBS, 200 µL of 4% PFA were added to each well and incubated for 1.5 h at RT. After two washes with PBS, the devices were stored at 4 °C until further processing.

#### Flow cytometry analysis of organoid pieces

At D120+30-40 of tract formation, organoid pieces were analysed by flow cytometry for Kaede tracing and pRabV tracing. For Kaede tracing, the two connected pieces of a tract were processes together; for pRabV tracing, only the contralateral organoid was processed. Organoids were dissociated in 150 µL of Trypsin (Thermo Fisher Scientific, 15400054)/Accutase (1:1), at 37 °C for 15 min, followed by manual pipetting with a P200 tip until a cell suspension without clumps was obtained. The action of Trypsin/Accutase was stopped by adding 400 µL of 0.1% BSA in PBS. The samples were analysed immediately after, in a NovoCyte Penteon Flow Cytometer. The data was analysed using the FlowJo Software

### Cryosectioning

For preparation of cryosections for immunofluorescent staining analysis, organoids (or organoid pieces) were fixed in 4% PFA for 1-2 hours at room temperature (RT). After three 15 min washes with PBS, organoids were immersed in a 15% Sucrose (Merck Millipore, 84097)/10% Gelatin (Sigma, G1890-500G) solution at 37 °C, until they sunk to the bottom of the tube (from 1h up to overnight). Organoids were embedded in the same sucrose/gelatin solution, solidified for 30 min at 4 °C, and flash frozen in a bath of 2- methylbutane super-cooled to a temperature of −50 °C with dry ice. Samples were stored at −70 °C until further processing. Cryoblocks were sectioned at 20 µm thickness using a cryostat (Thermo Fisher Scientific, CryoStar NX70).

### Immunofluorescent staining

#### Organoids

After the slides were defrosted and hydrated during 5 min in PBS, cryo-sections were permeabilized and blocked with blocking solution (5% bovine serum albumin (BSA; Europa Bioproducts, EQBAH-0500) and 0.3% Triton X-100 (Merck Millipore, 93420) in PBS) for 1-2 hours at RT. Antibody incubations were done in antibody solution (1% BSA, 0.1% Triton X-100 in PBS); after dilution of antibodies, the solution was spun at maximum speed for 2 min. First, sections were incubated with antibody solution containing 1:200 diluted primary antibodies, for 3.5 hours (up to overnight) at RT. Then, after one wash of 5 min with PBS, sections were incubated with antibody solution containing 1:500 diluted secondary antibodies and 1:10,000 diluted Hoechst 33342 nuclear dye (Thermo Fisher Scientific, H3569), for 1-2 hours at RT. Primary and secondary antibodies used in this study are summarized in **Table S2**. After one wash of 5 min with PBS, the slides were mounted with DAKO mounting medium (Agilent, S302380-2). Slides were left at RT to dry overnight and kept at 4 °C for long-term storage.

#### Axon tracts

Immunostaining of axon tracts was performed inside the microdevices, with the same solutions as described for organoid sections. To successfully stain the whole length of the tract, one of the two organoids was removed from the device and the solutions were passed through the channel carefully, by capillarity, from the side with an organoid towards the side without an organoid. Permeabilization/blocking was performed for 1.5 h, primary antibody exposure overnight, and secondary antibody exposure for 1 h. Two washes with PBS were done in between primary and secondary antibodies, and at the end. Stained tracts were stored at 4 °C, in PBS containing 0.01% of sodium azide.

#### Paraffin-embedded human brain samples

The use of human brain samples for histological analysis was approved by the local ethics committee of the Medical University of Vienna. Human brain material was processed for routine histopathology. The tissue was formalin fixed and embedded in paraffin and 3 µm thin FFPE tissue sections were prepared. Immunofluorescence for SATB2 was performed following the steps: slide dewaxing using Xylene Substitute (Epredia, 6764506); antigen retrieval (using sodium citrate buffer 10mM with 0.05% Tween, at pH 6: pre-heating at 100 °C; 30 min in a steamer; 15min cooling down at RT); two washes with TBS; one wash with TBS-T (TBS with 0.05% Tween); blocking buffer for 1 h at RT (5% BSA in TBS-T + 10% goat serum); primary antibody, 1:200 in 2% BSA TBS-T, overnight at 4 °C; wash with TBS-T; RbLMs linker (Anti-IgG1 + IgG2a + IgG3 antibody; Abcam, ab133469), 1:500 in 2% BSA TBS-T, 30 min at RT; secondary anti-Rabbit antibody (AF647), 1:500, 1 h at RT; wash with TBS-T; DAPI staining,10 min at RT in the dark; one wash with TBS-T; mounting with DAKO mounting medium; long-term storage at 4 °C.

### Western blotting

#### Sample preparation

Organoids from the same batch (typically 5) were collected and washed in PBS. All further steps were performed on ice. 300 µL of lysis buffer (RIPA with PhosStop and Proteinase Inhibitor, **Table S3**) was added. Organoids were dissociated by pipetting, until no clumps remained. 1 µL of Benzonase (Merck Millipore, US170664-3) was added and samples left on ice until an aqueous solution formed (30 min to 1 h). Samples were centrifuged at 15.000 g for 5 min and the supernatant transferred to a new tube.

#### Measurement of protein concentration

The Pierce Rapid Gold BCA protein assay kit (Thermo, 23227) was used to measure protein concentration, according to manufacturer’s instructions.

#### Gel running

Sample dilution was done using Laemmli Sample Buffer (Bio Rad, 161-0737), and incubated at 95°C for 5 minutes. 25 µg of protein was loaded per well. Samples were run on Nupage™ 3-8% Tris-Acetate Protein Gels (Thermo Scientific, EA03755BOX), using NuPAGE® Tris-Acetate SDS Running Buffer (20X, Thermo Fisher Scientific), at room temperature and 100 V. PageRuler™ Plus Prestained Protein Ladder, 10 to 250 kDa (Thermo Fisher Scientific, 26619) was used.

#### Gel transfer

Transfer to LiCOR membranes was done in NuPAGE® Transfer Buffer (Thermo Scientific, NP0006) with 10 % Methanol, for 1.5 h, at RT.

#### Blotting

After transfer, membranes were rinsed in PBS. Further steps were performed in 50 mL falcon tubes. Blocking was done with 5% BSA in PBS, 1 h at RT. Primary and secondary antibodies (**Table S2**) were diluted in PBS with 5% BSA and 0.1% Tween. Membranes were exposed to primary antibodies overnight at RT, and to secondary antibodies 1 h at RT. Membranes were washed in PBS with 0.1% Tween, 3 times 5 min in between antibodies and at the end. Finally, membranes were rinsed in water, and imaged.

### Image Acquisition

#### Cryosections

Immunostaining images were acquired with a Pannoramic FLASH 250 II digital Slide Scanner (3DHISTECH) at 20x magnification.

#### Live imaging of axon outgrowth

Axon outgrowth was imaged from 24 h to 72 h after the start of the tract formation assay, using a Celldiscoverer 7 (ZEISS). Acquisition parameters were as follows: objective 20x/0.7; 9 z-stacks with 2,5 µm steps; 16 Bits; 1x1 binning. Devices were fed after 2 days.

#### Widefield imaging of axon tracts

Axon tracts were imaged once a week (D8, D15, D22, D30), using a Celigo Image Cytometer (Nexcelom Bioscience).

#### Confocal imaging of stained axon tracts

Immunostaining of axon tracts was performed in an inverted Scanning confocal microscope LSM800 (Zeiss), with a 20x objective.

#### Western blots

Western blot images were acquired using a ChemiDoc MP Imaging System (Bio-Rad).

### Imaging data analysis

All data matrices of quantifications were processed in R software v4.2.2 using dplyr (v1.0.9) and visualized using ggplot2 (v3.4.0). Boxplots represent the median value; two hinges corresponding to the first and third quartiles (the 25th and 75th percentiles); and whiskers extending from the hinge to the highest/lowest value no further than 1.5 * IQR from the hinge (where IQR is the inter-quartile range, or distance between the first and third quartiles). Statistical analyses were performed in R software (**Table S4**).

#### Organoid area

Measurement of organoid area was performed manually by drawing regions of interest (ROIs) in the CaseViewer software (3DHISTECH), using the largest cross-section per organoid. Plotting was done with the geom_smooth() function of ggplot2, using method “lm” and formula y ∼ x + I(x^2^). Number of organoids included per genotype and timepoint: D40, 13 *ARID1B*^+/+^, 9 *ARID1B*^+/-^; D60, 22 *ARID1B*^+/+^, 37 *ARID1B*^+/-^; D90, 26 *ARID1B*^+/+^, 58 *ARID1B*^+/-^; D120, 19 *ARID1B*^+/+^, 25 *ARID1B*^+/-^.

#### Western blot band intensities

Measurement of band intensity for each staining was done by manually drawing equally sized rectangles around the bands, using the “Rectangle” tool in Fiji. The band intensity of mSWI/SNF proteins was normalized to the intensity of the respective β-Actin band. The mean of *ARID1B*^+/+^ normalized values was considered 1. The fold-change between normalized intensities was plotted.

#### Density of SATB2^+^ cells

For detecting positive nuclei, a custom workflow was designed, using ”CaseViewer”, ”Fiji” and ”Cellpose”. The organoid slices stained against SATB2 were scanned, marked, and exported as individual channels via the CaseViewer software. In Fiji, we performed segmentation of the outer-most organoid surface, where SATB2^+^ neurons are abundant, with a 325μm-thick band (1000 pixels), using the “Segmented line” tool. To map this region to an xy-coordinate system, reduce image size and exclude unwanted areas the ”Straighten” command was used. Via the Cellpose implementation in Fiji, the positive cells in each channel were detected using the pre-trained ”Cyto” model. Positive cells were segmented, and regions of interest generated. Exemplary outputs of the different steps of the protocol are depicted in **Fig.S4C**.

#### Axon outgrowth at 72 h

For analyzing the number of axons extending between the two organoids we designed a custom script written in Fiji-Macro language. In addition to the built-in tools, we used the MorpholibJ collection (https://imagej.net/plugins/morpholibj) as well as the “Derivatives” filter from the FeatureJ collection of the ImageScience update site (https://imagej.net/libs/imagescience). The images of axon outgrowth were acquired in 3D using oblique illumination (9 z-stacks with 2.5µm steps). For extracting the axons, the derivative image in z-direction was computed, which significantly enhanced the contrast. Then, the slice of highest contrast (maximum standard deviation) was chosen for further analysis. We needed to restrict the analysis to the inside of the microdevice channel. Therefore, we determined its upper and lower border using the “Tubeness” filter on a max-projection and then thresholding the result. Left and right borders were found via plotting intensities along x and finding the two maxima. Line ROIs for the cross-section measurements were created within the area of the chamber in a sampling distance of 50 pixels (32.5 µm). For the final axon detection, we used the derivative filtered image, applying the “Directional Filtering” from MorpholibJ as well as the “Tubeness” filter to enhance the contrast of the axons. At the end, the individual line-ROIs were applied, and intensities plotted. Maxima above a certain threshold (2000) along the line were considered as crossing axons. Exemplary outputs are depicted in **Fig.S10**. The number of axons per sampling line was counted throughout the length of the chamber. For data visualization, the average number of axons in the first five bins (162.5 µm) and along the first 35 bins (1.1 mm) was computed and plotted.

#### Axon outgrowth speed

Axon outgrowth speed was calculated from manual tracking of growth cones, using the “MtrackJ” tool (*152*) in Fiji.

#### Axon tract thickness

Average axon tract thickness was calculated based on seven equidistant measurements along the axon tract, marked manually using the “Line” tool in Fiji.

#### Co-localization of Crimson and SATB2 in Rabies virus experiments

Co-localization of Crimson and SATB2 was done by visual assessment of immunofluorescence stainings. All Crimson^+^ cells present in a certain organoid section were quantified for SATB2 positivity/negativity, and the percentage of SATB2^+^ cells plotted.

### Single-cell RNA sequencing

#### Generation of single-cell suspension of organoid cells

Organoids used for scRNAseq were harvested, cut into 2 or 3 pieces using two P10 pipette tips, and washed with PBS. Each individual organoid was incubated in 1.5 mL of Trypsin/Accutase (1:1) containing 1 µL/mL of TURBO™ DNase (Thermo Fisher Scientific, AM2238, 2 U/µL) in a gentleMACS Dissociator (Miltenyi Biotec, 130-093-235) in the program NTDK1. After dissociation, tubes were briefly spun down and 1.5 mL of buffer (ice-cold PBS with 0.1% BSA) was added to the dissociated cells. The cell suspension was spun at 400 g for 5 min at 4 °C. The supernatant was aspirated, leaving a margin of around 200 µL, and cells were resuspended in 700 µL of buffer. The suspension was filtered through a 70 µm strainer once, and through a FACS tube cap twice.

#### Organoid multiplexing

The 10X Genomics 3’ CellPlex Kit was used to multiplex each individual organoid with a unique Cell Multiplexing Oligo (CMO), as described in the manufacturer’s protocol, except for the use of the aforementioned buffer formulation and 400 g in the spinning steps.

#### FACS sorting of viable cells

After multiplexing, the viability dye DraQ7 (Biostatus, DR70250, 0.3 mM) was added at a concentration of 20 µL/mL and the suspension gently mixed with a P1000 pipette. 50k live cells of each individual organoid were FACS sorted onto the same tube using a 100 µm nozzle; singlets were gated based on forward and side scatter, and live cells based on negative excitation with an Alexa 700 filter.

#### Library preparation

The final pool of multiplexed cells was spun and resuspended in a small volume, and live cells were counted again. Libraries were loaded with 50k live cells per channel (to give estimated recovery of 15k cells per channel) onto a Chromium Single Cell 3′ B Chip (10X Genomics, PN-1000073) and processed through the Chromium controller to generate single-cell GEMs (Gel Beads in Emulsion). ScRNAseq libraries were prepared with the Chromium Single Cell 3′ Library & Gel Bead Kit v.3 (10x Genomics, PN-1000075).

#### Library sequencing

Libraries were pair-end sequenced in a NovaSeq S4 flowcell.

### Single-cell RNA sequencing data analysis

#### Dataset information

Four different sequencing experiments were performed in four different libraries. Each library contained organoids at D120 (±5 days), from the same batch (EBs generated on the same date), of the following cell lines and clones:

1. Library 177136: *ARID1B*^+/+^ clone 2c (2 organoids), *ARID1B*^+/-^ clone 2a (2 organoids).
2. Library 178119: *ARID1B*^+/+^ clone 2c (3 organoids), *ARID1B*^+/-^ clone 2a (3 organoids).
3. Library 178120: *ARID1B*^+/+^ clone 2d (3 organoids), *ARID1B*^+/-^ clone 2b (3 organoids).
4. Library 184337: *ARID1B*^+/-^ clone 1a (3 organoids), *ARID1B*^+/+^ clone 3a (3 organoids), *ARID1B*^+/-^ clone 3b (3 organoids).

#### Data pre-processing

ScRNA-seq reads were processed with Cell Ranger multi v6.1.1 using the prebuilt 10X GRCh38 reference refdata-gex-GRCh38-2020-A, and including introns. Further processing such as dimensionality reduction, clustering, and visualization of the scRNA-seq data was performed in R v4.2.2 with Seurat v4.3.0.

#### Identification of non-telencephalic clusters

Non- telencephalic cells were separated at clustering resolution 0.02 (“FindClusters”). One non-telencephalic cluster (cluster 2) was removed for downstream analysis, based on absent/low FOXG1 expression. A total of 1.76% of cells were excluded by this filter.

#### Removal of low-quality cells

After exclusion of non-telencephalic cells, genes located on the Y-chromosome and those not detected in at least 5 cells were excluded. We used the Gruffi algorithm to identify and remove cells presenting a transcriptional signature of cellular stress (*153*). Cells were removed if they were identified as stressed, the gene number metric was below a MAD (median absolute deviation) based threshold (library: 178119 <900, 178120 <1100, 184337 <1200, 177136 <1500), the percentage of mitochondrial reads was 10% or above, and the percentage ribosomal RNAs was 25% or above. Count data was log-normalized and scaled. Dimensionality reduction was performed using PCA on the top 3000 most variable genes. Batch correction across libraries was performed using Harmony v0.1.1, and the 40 harmony embeddings were used for UMAP visualizations.

#### Identification of telencephalic clusters

The first 10 Harmony dimensions were used to identify neuronal subclusters with a resolution of 0.2. Clusters were assigned to cell types using known neuron markers and the developing human neocortex reference (PMID: 31303374): INs (cluster 5), dividing progenitors (cluster 3), RGs (cluster 4), IPCs (cluster 2), ExN1 (cluster 0), ExN2 (cluster 1), and ExN3 (cluster 6).

#### Differential gene expression analysis

Wilcoxon rank sum test implemented in Presto v1.0.0 was used to identify differentially expressed genes between *ARID1B*^+/+^ and *ARID1B*^+/-^ on the subset of SATB2^+^ cells for each sequencing library. ClusterProfiler v4.6.2 and enrichR v3.1 were used for further functional analysis and visualization thereof, using the top 300 dowregulated genes in *ARID1B*^+/-^ organoids, per pairwise comparison. The most statistically significant terms were considered as those found in 5 out of 5 comparisons, with a gene ratio above 5%, and ordered by the average P value (Pval) across comparisons.

#### Data plotting

Two-dimensional representations were generated using uniform manifold approximation and projection (UMAP) (uwot v0.1.14). Visual representation of differentially expressed genes was generated using ggplot2 and cnetplot. Data matrices of quantifications were processed in R software v4.2.2 using dplyr (v1.0.9) and visualized using ggplot2 (v3.4.0).

---

**Figure S1. (Related to Fig.1B-C and G).**
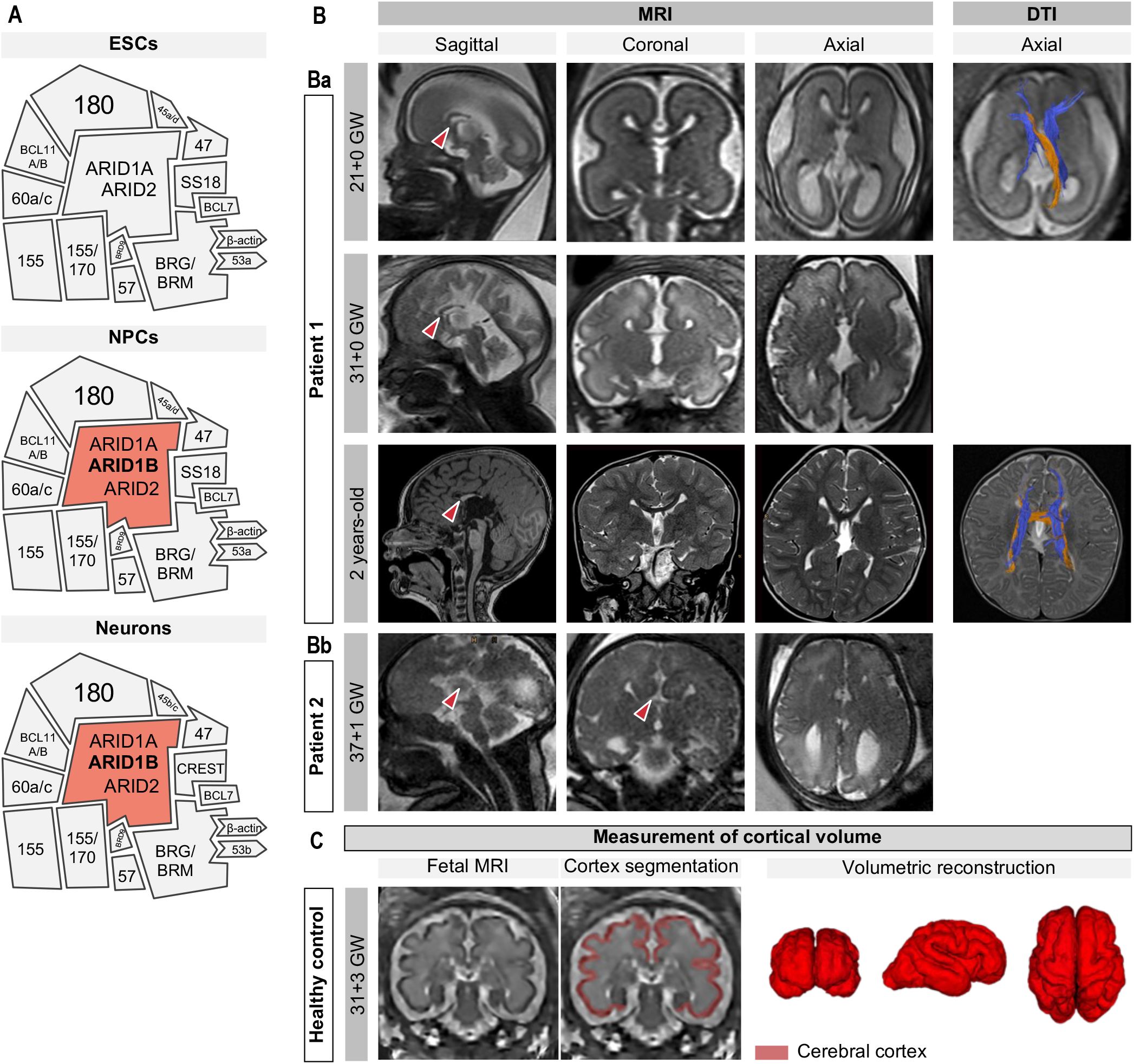
Recruited ARID1B patients were diagnosed with corpus callosum malformations during gestation. **(A)** Distinct mSWI/SNF configurations that exist during development. ARID1B is absent from the complex in embryonic stem cells (ESCs); and is a core member in neural progenitor cells (NPCs) and neurons, interchangeable and mutually exclusive with ARID1A and ARID2. The numbers indicate BAF subunits (e.g.: 150, BAF150). Adapted from (*7*). **(B)** Fetal and postnatal magnetic resonance imaging (MRI) and diffusion tenson imaging (DTI) examinations of recruited patients harboring *ARID1B* mutations. Sagittal, coronal, and axial imaging planes are shown. **(Ba)** Patient 1 was examined at 21+0 gestational weeks (GW) (T2-weighted images), at 31+0 GW (T2-weighted images), and postnatally at 2 years (coronal and axial T2-weighted; sagittal T1 weighted images). Note the short and truncated corpus callosum in the sagittal planes (red arrowheads). DTI at 21+0 GW prenatally and 2 years postnatally show Probst bundles (blue), large bilateral, barrel-shaped axonal structures that form longitudinal intrahemispheric bundles situated in both hemispheres (*154*); and sigmoid bundles (orange), ectopic tracts crossing from left occipital lobe to right frontal lobe, and vice versa (*155*). **(Bb)** Prenatal MRI of Patient 2 at 37+1 GW. Note the complete absence of the corpus callosum (red arrowheads). Images from both patients reveal age-appropriate cortical maturation and normal gyrification. **(C)** Fetal MRI based super-resolution reconstruction and cortical plate volume measurement; example of a healthy control at 31+3 GW. Atlas-based segmentation (*138*) of the cortex was performed using the open-source application ITK-SNAP, allowing volumetric reconstruction and measurement of cortical volume in patients and healthy control individuals.

**Figure S2. (Related to Fig.1C-D).**
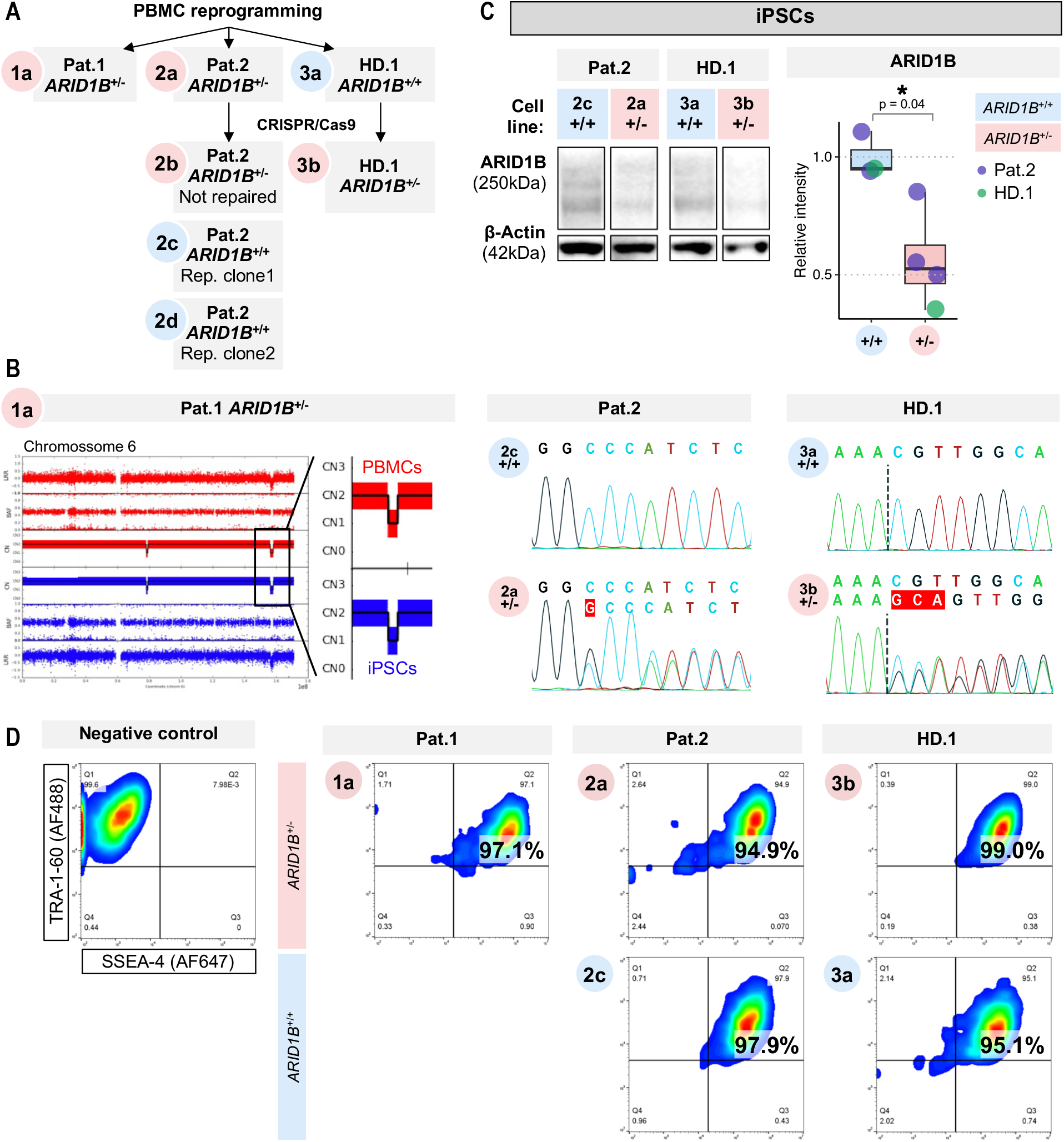
Characterization of patient iPSCs reveals *ARID1B* mutations and reduced ARID1B protein levels. **(A)** Reprogrammed and engineered cell lines used in this study. IPSCs were obtained from Patient 1 (Pat.1), Patient 2 (Pat.2), and Healthy Donor 1 (HD.1) by reprogramming of peripheral blood mononucleated cells (PBMCs). CRISPR/Cas9 genome editing was used to generate isogenic repair *ARID1B*^+/+^ cell lines from Pat.2 iPSCs and *ARID1B*^+/-^ mutant cell lines from HD.1 iPSCs. For Pat.2, a clone that went through the CRISPR/Cas9 genome editing procedure but did not get edited was used in scRNAseq experiments to control for off-target effects (clone 2b). **(B)** Confirmation of *ARID1B* gene sequence in patient and control cell lines. Single nucleotide polymorphism (SNP) genotyping of Pat.1 PBMCs and iPSCs shows a heterozygous microdeletion (copy number CN = 1) in the region q25 of chromosome 6, encompassing *ARID1B*. Sanger sequencing of Pat.2 and HD.1 iPSCs shows wild-type sequence in *ARID1B*^+/+^ cell lines and heterozygous frame shift mutations by nucleotide insertions in *ARID1B*^+/-^ cell lines (red nucleotides: Pat.2, NM_020732.3: c.2201dupG; HD.1, NM_020732.3: c.3489_3490insNN). **(C)** Western blot analysis of ARID1B protein levels in iPSCs shows a two-fold reduction in *ARID1B*^+/-^ cell lines, compared to isogenic controls. The boxplot shows the median value; each datapoint is an individual sample (iPSCs pooled from the same culturing well); datapoint colors indicate genetic background. Statistical test is analysis of variance (ANOVA); 0 ≤ p < 0.001, ***; 0.001 ≤ p < 0.01, **; 0.01 ≤ p < 0.05, *; p ≥ 0.05, ns (see results of statistical tests in **Table S4**). **(D)** Immunostaining and flow cytometry analysis, shown as density plots, of pluripotency markers TRA- 1-60 (conjugated with AF488) and SSEA-4 (AF647); gating is based on negative control (unstained sample). Note that 94% or more cells are double-positive in all cell lines.

**Figure S3. (Related to Fig.1C-D).**
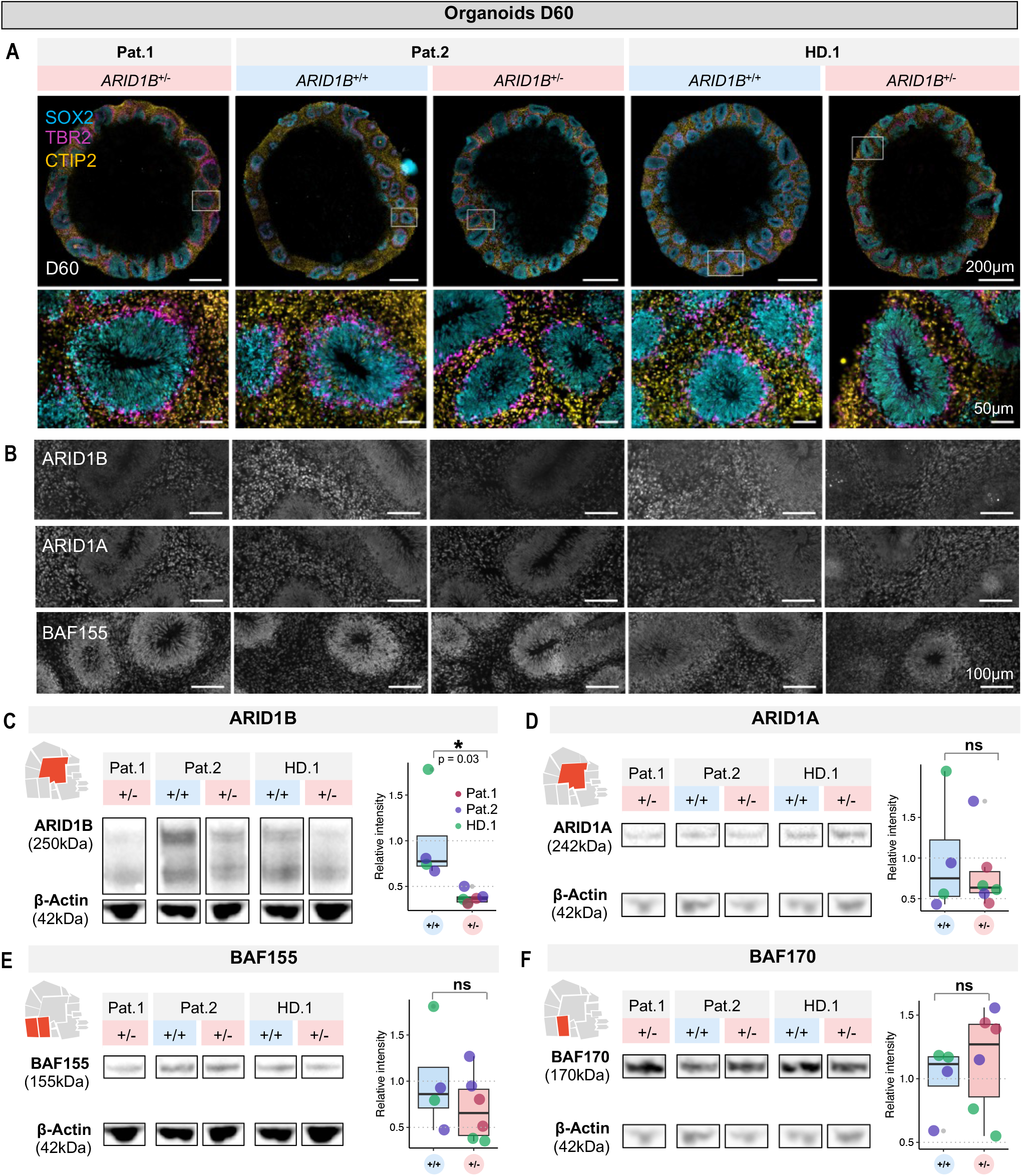
Reduced levels of ARID1B do not affect the differentiation of dorsal cortical progenitors and early-born neurons in organoids. **(A)** Immunostaining analysis of dorsal cortical specification markers in organoids at D60. Neural rosettes with stratified organization of SOX2^+^ radial glia cells, TBR2^+^ intermediate progenitor cells, and CTIP2^+^ early-born excitatory neurons are formed in *ARID1B*^+/+^ and *ARID1B*^+/-^ organoids. **(B)** Immunostaining of ARID1B, ARID1A, and BAF155 shows reduced signal of ARID1B and unaffected signal of ARID1A or BAF155 in progenitor and neuron zones of *ARID1B*^+/-^ organoids, compared to *ARID1B*^+/-^ organoids. **(C-F)** Western blot analysis of ARID1B **(C)** shows reduction of protein levels in D60 *ARID1B*^+/-^ organoids; ARID1A **(D)**, BAF155 **(E)**, and BAF170 **(F)** show comparable levels between *ARID1B*^+/-^ and *ARID1B*^+/+^ D60 organoids. In western blot analyses, the boxplots show the median value; each datapoint is an individual sample (5 organoids pooled from the same batch); datapoint colors indicate genetic background. Statistical tests are analysis of variance (ANOVA); 0 ≤ p < 0.001, ***; 0.001 ≤ p < 0.01, **; 0.01 ≤ p < 0.05, *; p ≥ 0.05, ns (see results of statistical tests in **Table S4**).

**Figure S4. (Related to Fig.1F).**
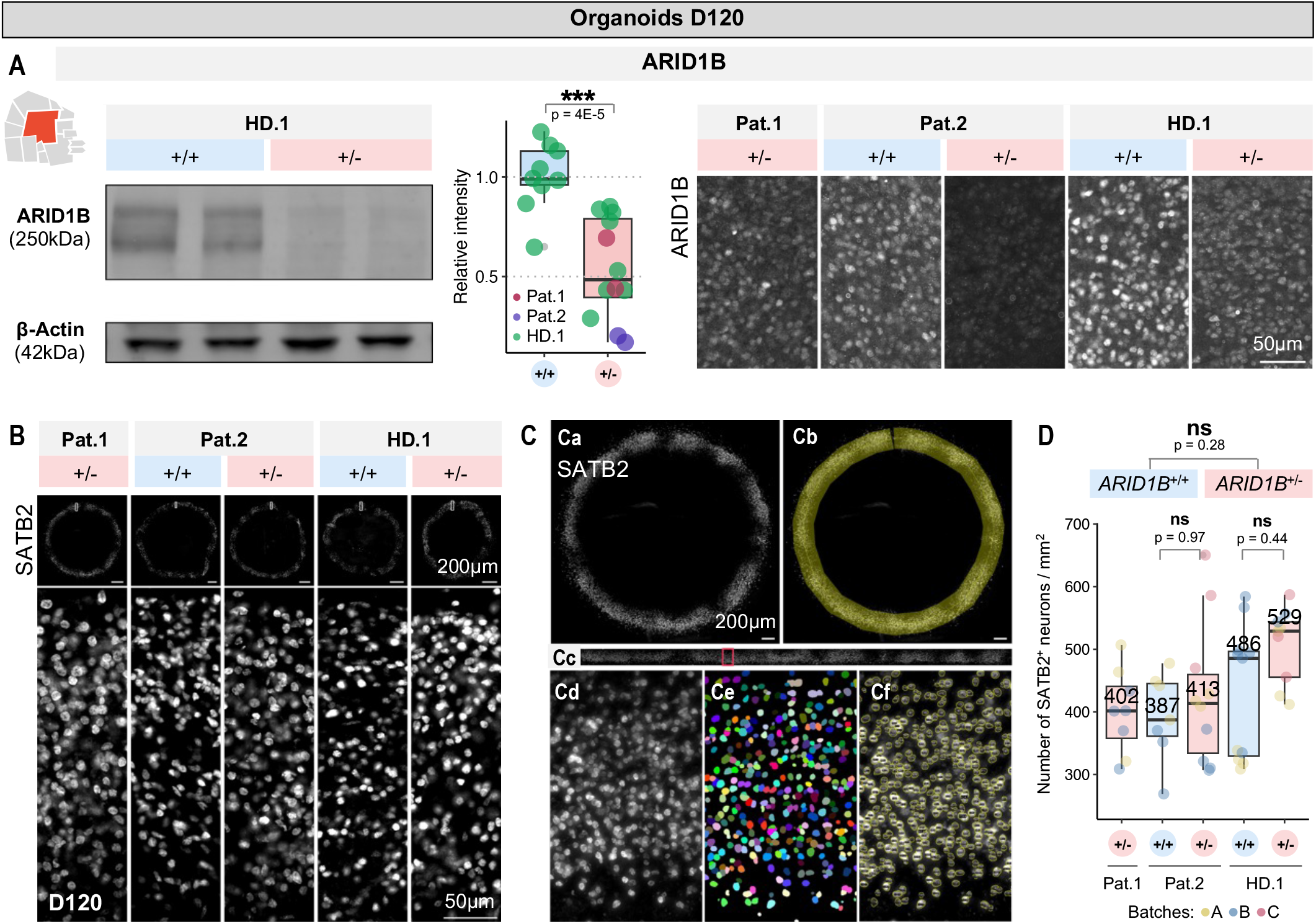
Lowered ARID1B levels do not affect the differentiation of SATB2^+^ neurons in *ARID1B*^+/-^ organoids. **(A)** Western blot and immunostaining analysis of ARID1B levels at late stages of organoid development shows reduction of protein levels in D120 *ARID1B*^+/-^ organoids. In western blot analysis, the boxplots show the median value; each datapoint is an individual sample (5 organoids pooled from the same batch); datapoint colors indicate genetic background. In immunostainings, organoid sections of all cell lines were stained simultaneously, imaged with the same settings, and without any post-processing adjustment. **(B)** Immunostaining of SATB2 in organoids of all cell lines at D120. At this stage, *ARID1B*^+/-^ and *ARID1B*^+/+^ organoids present abundant SATB2^+^ neurons lining their outer-most surface. **(C)** Method for segmentation of SATB2^+^ neurons: **(Ca)** organoid section stained against SATB2 (nuclear staining); **(Cb)** segmentation of the outer- most organoid surface, where SATB2^+^ neurons are abundant, with a 325µm-thick band (1000 pixels, “Segmented line” tool in FIJI); **(Cc)** straightening of the segmentation (”Straighten” tool in FIJI); **(Cd-Cf)** segmentation of individual cells and generation of regions of interest (details in **Materials and Methods**). **(D)** Quantification of the number of SATB2^+^ neurons per mm^2^. The boxplots show the median value; each datapoint is an individual organoid (technical replicate); datapoint colors indicate organoid batches (biological replicates). Statistical tests are analysis of variance (ANOVA); 0 ≤ p < 0.001, ***; 0.001 ≤ p < 0.01, **; 0.01 ≤ p < 0.05, *; p ≥ 0.05, ns (see results of statistical tests in **Table S4**).

**Figure S5. (Related to Fig.2A-D).**
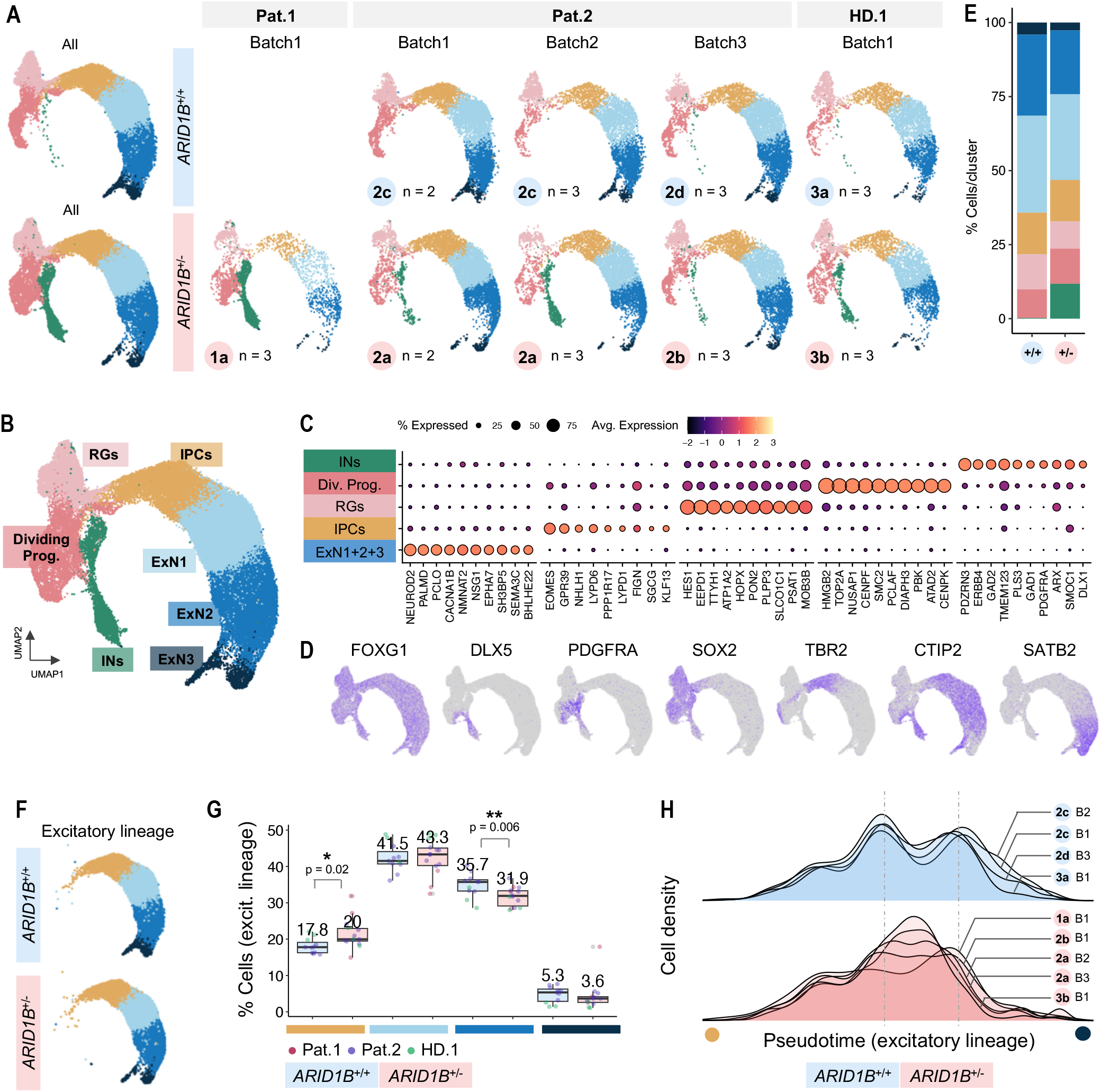
ScRNAseq analysis allows the recovery of telencephalic subpopulations and reveals changes in excitatory lineage trajectories. **(A)** UMAP projection per genotype, cell line, and batch. A total of 11 control and 14 patient organoids were tagged with unique molecular identifiers and could be demultiplexed for downstream analyses. Cell line clone identifiers (1a, 2a, 2b…) are as in Fig.2A. **(B)** UMAP projection and unbiased clustering at resolution 0.2 shows seven clusters of telencephalic populations: interneurons (INs), dividing progenitors, radial glia (RGs), intermediate progenitor cells (IPCs), and three populations of increasingly mature deep- and upper-layer excitatory neurons (ExN1, ExN2, ExN3). **(C)** Top 10 markers per cluster (using the “FindMarkers” tool of Seurat). **(D)** Cell type-specific marker genes commonly used to identify telencephalon (FOXG1), INs (DLX5), oligodendrocyte precursor cells (PDGFRA), neural progenitors (SOX2), deep-layer ExNs (CTIP2) and upper-layer ExNs (SATB2). **(E)** Percentage of cells per cluster in *ARID1B*^+/+^ and *ARID1B*^+/-^ datasets. **(F)** UMAP projection of clusters of the dorsal excitatory lineage (•IPCs, •ExN1, •ExN2, •ExN3). **(G)** Percentage of cells per cluster. Median values are indicated; each datapoint is a single organoid; colors mark different genetic backgrounds. Statistical tests are analysis of variance (ANOVA); 0 ≤ p < 0.001, ***; 0.001 ≤ p < 0.01, **; 0.01 ≤ p < 0.05, *; p ≥ 0.05, ns (see results of statistical tests in **Table S4**). **(H)** Slingshot analysis for pseudotime visualization of cell density along the excitatory lineage (from •IPCs to •ExN3), per individual clone and batch; each line is the average of 2 or 3 organoids (as in **(A)**).

**Figure S6. (Related to Fig.2E-G).**
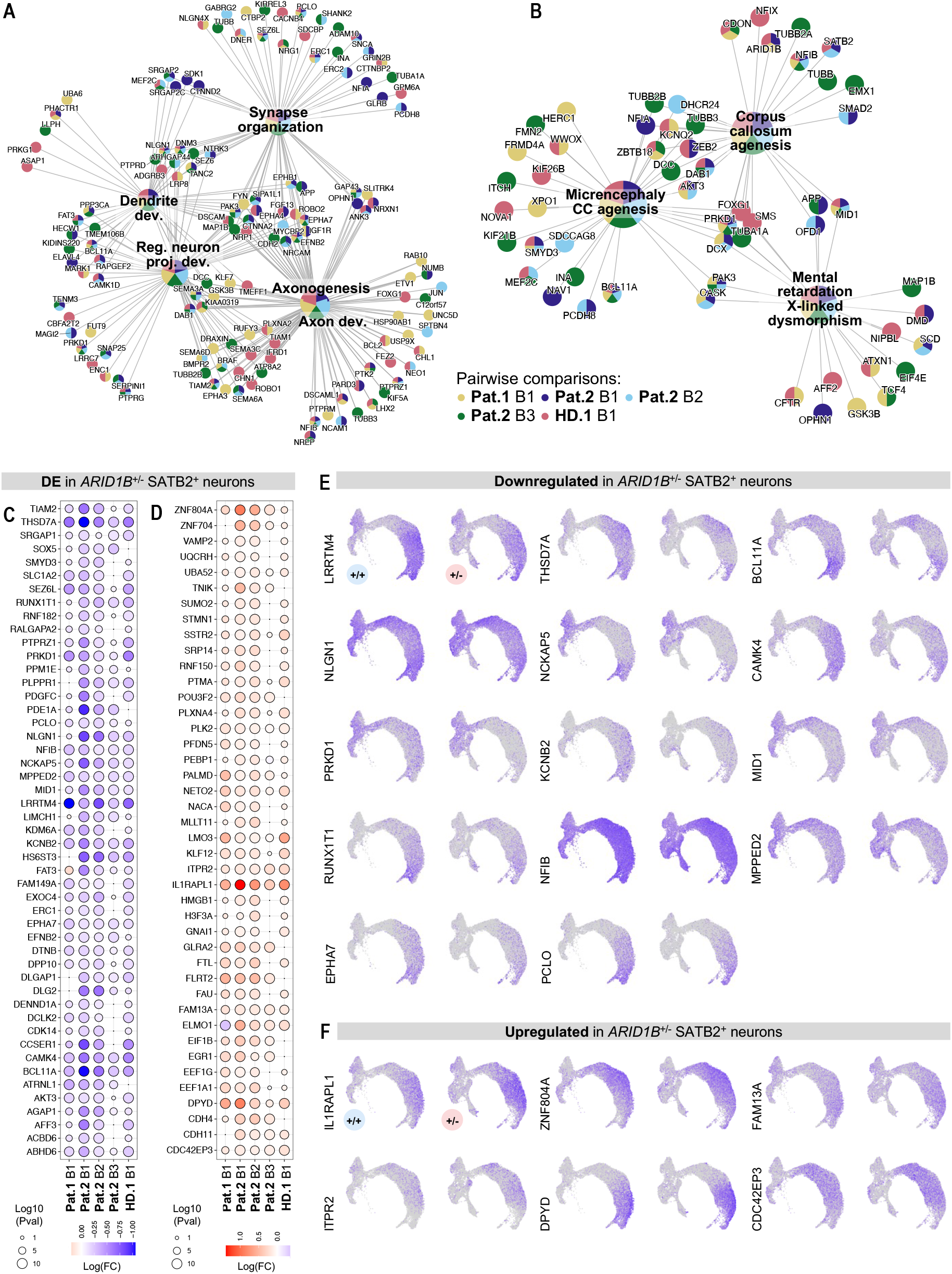
Differentially expressed genes in SATB2^+^ neurons of *ARID1B*^+/-^ organoids are related to ID and ACC. **(A-B)** Individual genes found associated with top gene ontology (GO) terms of cell processes **(A)** and rare diseases **(B)**, using the top 300 dowregulated genes in *ARID1B*^+/-^ organoids. Colors indicate the pairwise comparisons in which each gene was found downregulated. **(C-D)** Up- **(B)** and downregulated **(D)** Genes in *ARID1B*^+/-^ SATB2^+^ ExNs found differentially expressed in 4 out of 5 or 5 out of 5 pairwise comparisons, ordered alphabetically (violet, downregulated; red, upregulated). **(E-F)** Expression of DE genes found up- **(E)** and downregulated **(F)** in 5 out of 5 pairwise comparisons (top-ranked candidates shown in Fig.2G).

**Figure S7. (Related to Fig.3A).**
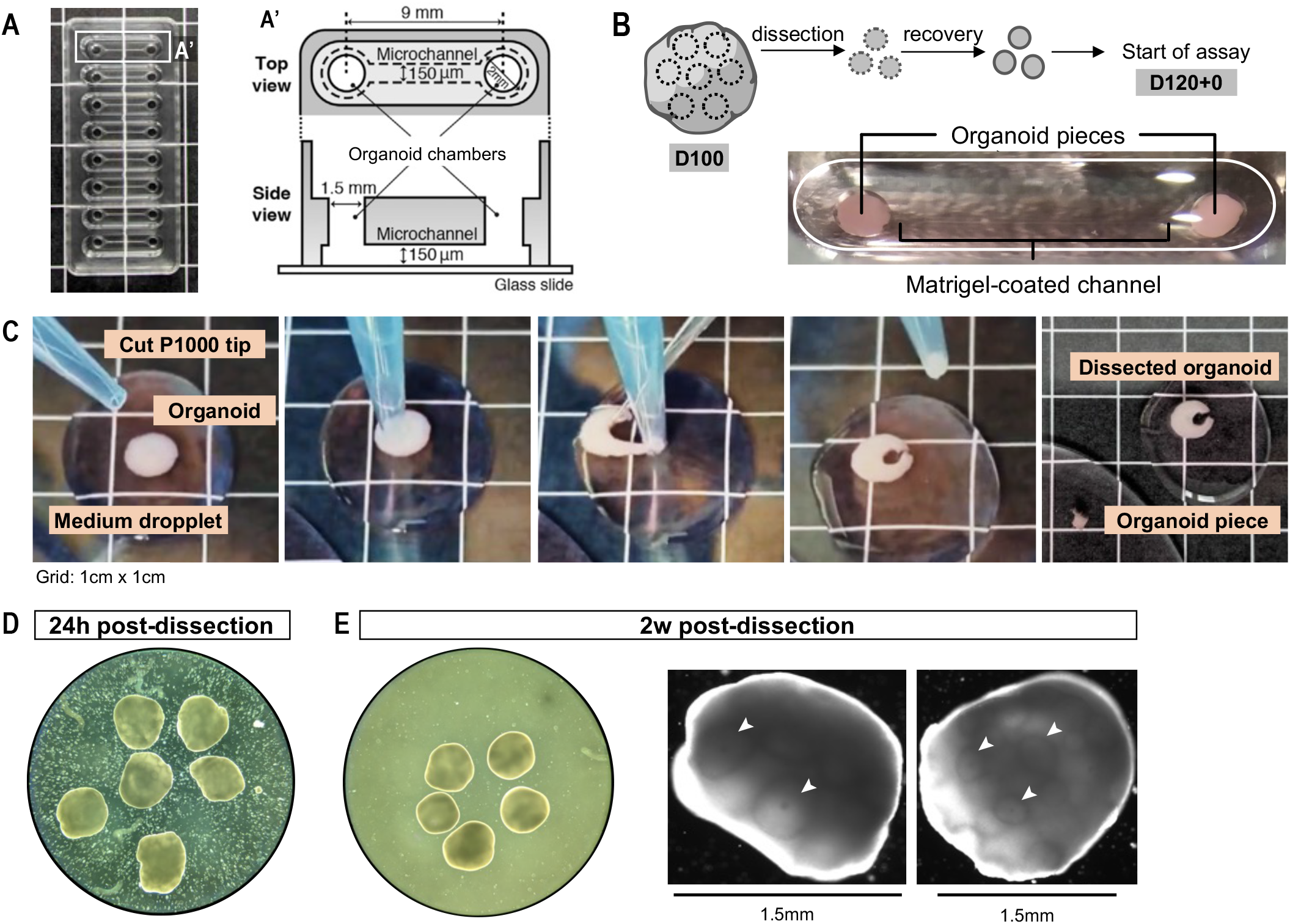
Tract formation from dissected organoids occurs in bioengineered microdevices. **(A)** Microdevices used for axon tract formation assay. **(B)** Schematic representation of the protocol. Organoids were dissected at D100 and placed in two wells interconnected by a Matrigel-coated channel at D120. This marks t = 0 (D120+0) of the assay. **(C)** Dissection protocol. Each organoid is used to obtain 5 to 10 dissected pieces. **(D-E)** Organoid pieces 24 h **(D)** and 2 weeks **(E)** after dissection. Note the appropriate size (1-1.5mm) and healthy appearance of the tissue after two weeks, with minimal cell death and visible rosette structures (arrowheads, ➤).

**Figure S8. (Related to Fig.3B-C and H-I).**
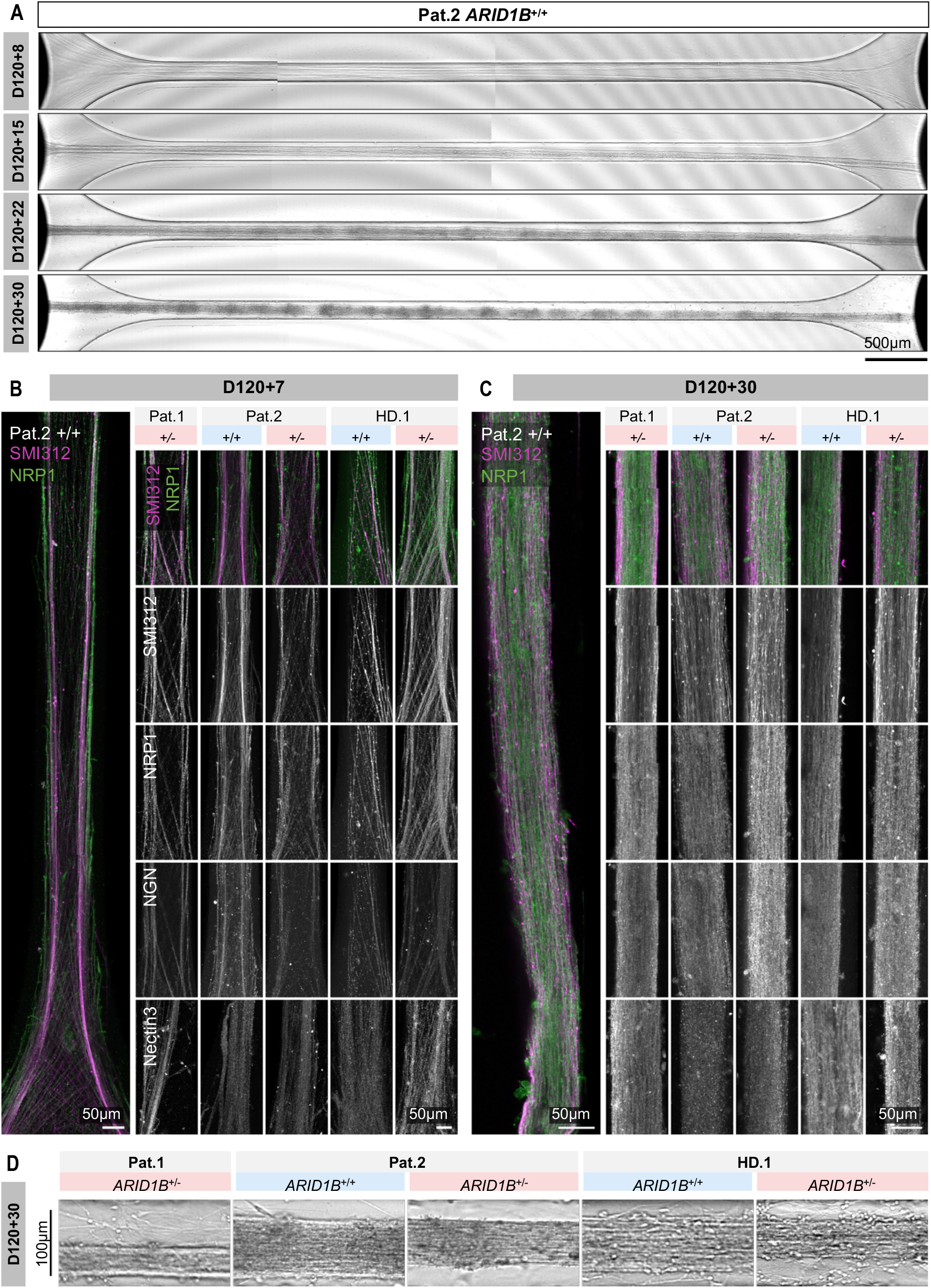
The thickness of *ARID1B*^+/-^ CC-like axon tracts from late-stage organoids is reduced. **(A)** Example of tract formation timepoints (D120+8/15/22/30). Note that at D120+8 several thin bundles from opposing organoids have already met at the middle of the channel. These different bundles come together by D120+15, to form a single tract that increases in thickness until D120+30. **(B-C)** Representative images of immunostaining analysis of axonal markers NFH, NRP1, Nectin3, and Neogenin, at D120+7 **(B)** and D120+30 **(C)** of axon tract formation. Each staining was verified in 2 or 3 different batches (1 tract/batch) of all 5 cell lines. **(D)** Representative images at D120+30 showing reduced thickness of *ARID1B*^+/-^ tracts.

**Figure S9. (Related to Fig.3D-G and J-K).**
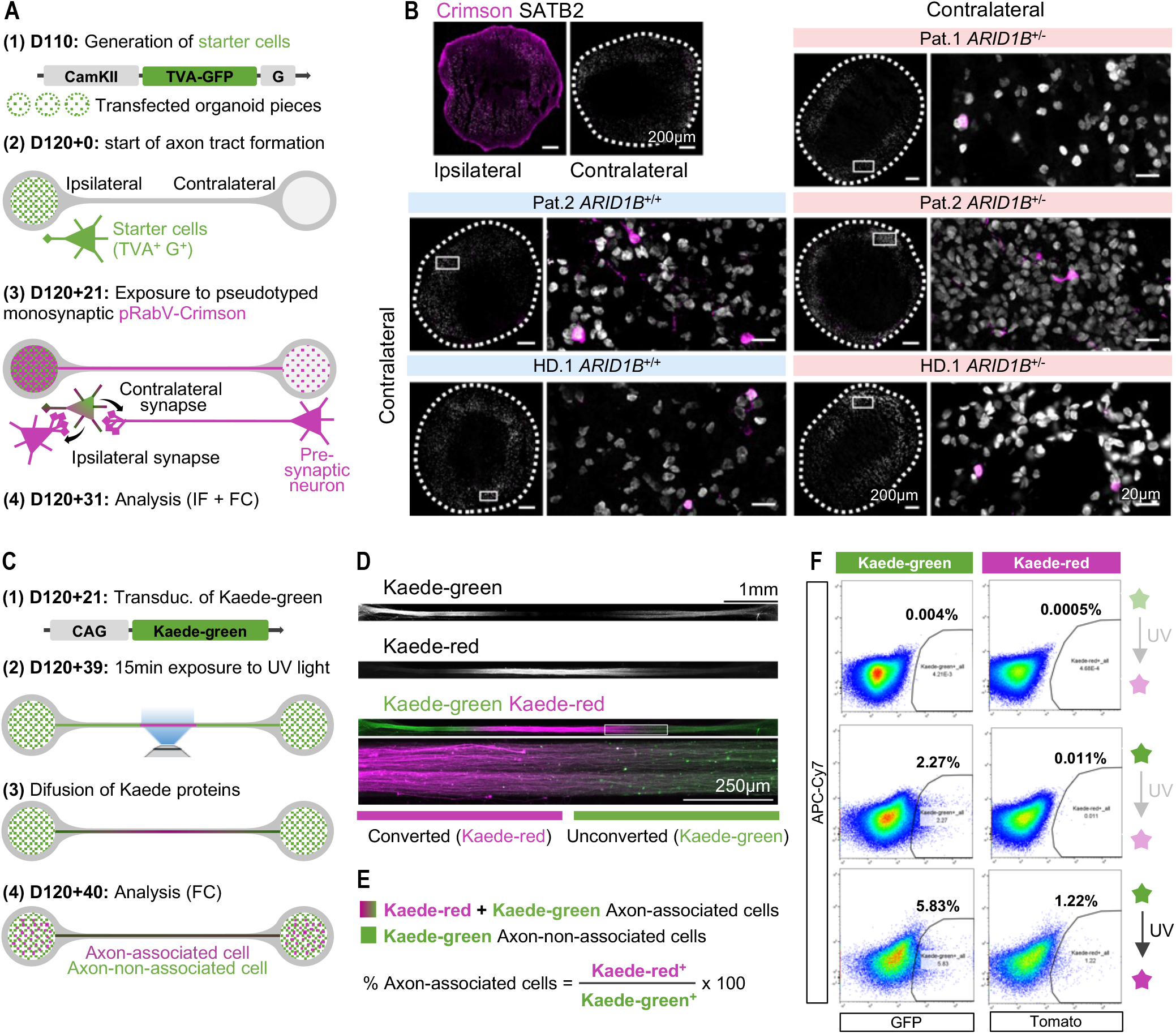
Rabies virus tracing and Kaede protein photoconversion can be used to assess tract connectivity. **(A)** Schematic representation of the protocol used to assess the number and cell-type identity of contralaterally connected cells. 1) Expression of CamKII::TVA-GFP-G was achieved by adeno-associated virus (AAV)-mediated transfection of organoid pieces at D110 (before start of axon tract formation), generating GFP^+^ starter cells (SCs). 2) For the axon tract formation assay, only one of the two organoid pieces contained SCs, ensuring unilateral pRabV entry. 3) Connected organoids were exposed to Crimson-expressing pRabV at D120+21; pRabV entry originated GFP^+^Crimson^+^ SCs, and pRabV spread originated GFP^-^Crimson^+^ pre-synaptic connected cells. Connected cells were ipsilateral, when on the same organoid as the SC population; or contralateral, when spread happened through the axon tract. 4) 10 days after pRabV exposure, at D120+31, organoids were analysed via immunofluorescent staining (IF) or flow cytometry (FC). **(B)** Co-immunostaining of Crimson and SATB2, showing abundant ipsilateral and sparse contralateral Crimson^+^ cells. Co-localization of Crimson and SATB2 allowed the identification and quantification of SATB2^+^ callosal projection neurons performing contralateral connections in *ARID1B*^+/+^ and *ARID1B*^+/-^ organoids of all cell lines. **(C)** Schematic representation of the protocol used to assess the proportion of axon-associated cells. 1) Expression of CAG::Kaede-green was achieved by AAV-mediated transfection at D120+21 days, resulting in mosaic expression in organoid cells, including projection neurons, and subsequent diffusion to the axon tract. 2) At D120+39, the central region of the axon tract was exposed to UV light during 15 min, leading to local conversion of Kaede-green into Kaede-red. 3) Kaede-red subsequently diffused to the cell bodies of axon-associated cells, 4) allowing their identification with flow cytometry (FC) analysis 24 h later. **(D)** Live imaging of Kaede-green and Kaede-red immediately after photo- conversion. Note that the conversion is confined to the central tract region (Kaede-red^+^). **(E)** 24 h after photo-conversion, axon-associated cells show green and red fluorescence; axon-non-associated cells show just green fluorescence. **(F)** Exemplary plots of flow cytometry analysis: negative controls with no Kaede transfection **(top)** or no photoconversion **(middle)**; and sample with local Kaede photoconversion in the axon tract **(bottom)**.

**Figure S10. (Related to Fig.4C-E).**
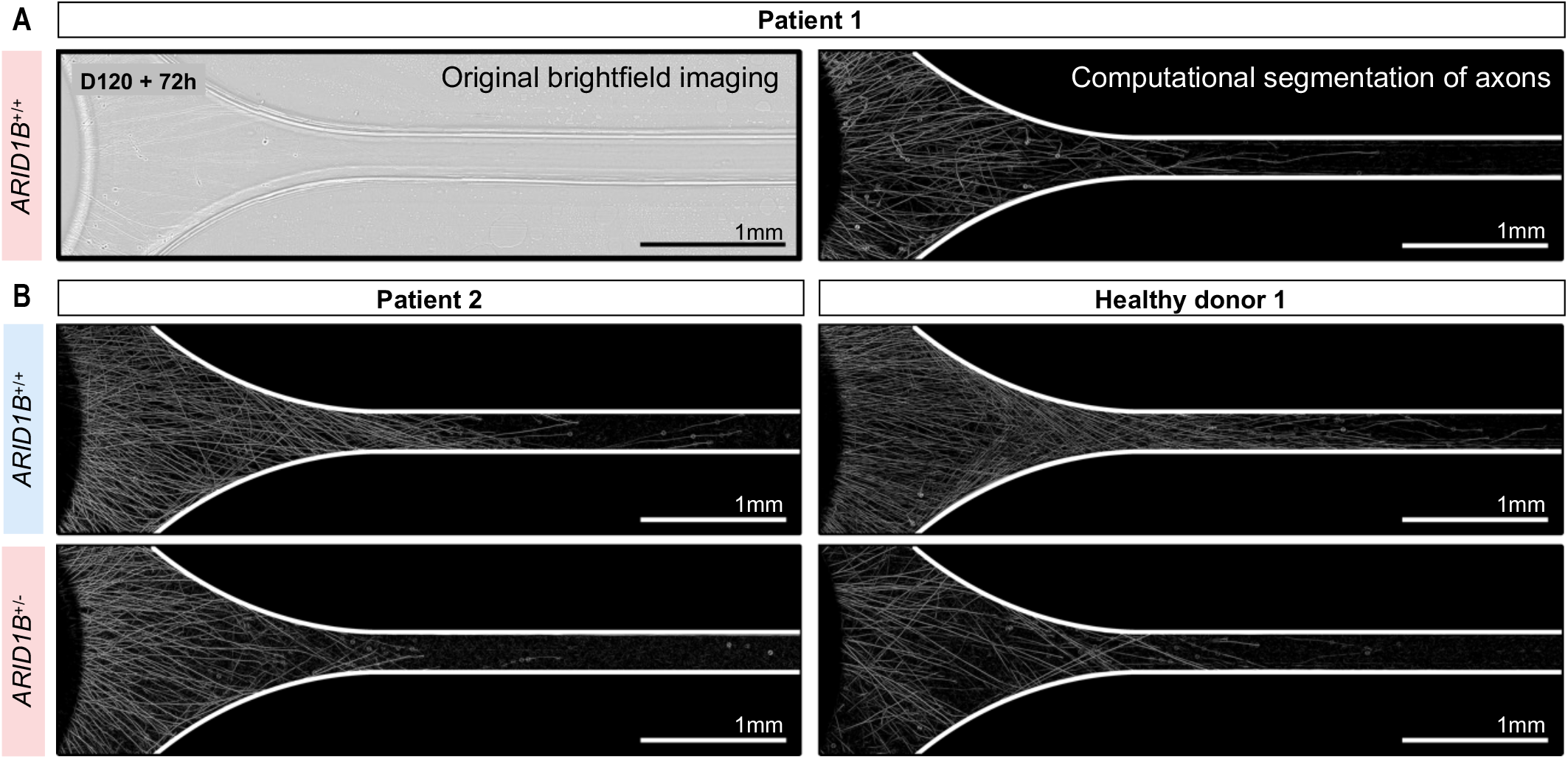
*ARID1B*^+/-^ organoids show decreased axon outgrowth at D120+72h. **(A)** Automatized analysis pipeline of images acquired with oblique illumination and 9 z-stacks **(left)**, used for axon segmentation and quantification **(right)** (details in **Materials and Methods**). **(B)** Representative images showing reduced axon outgrowth in *ARID1B*^+/-^ organoids.

**Figure S11. (Related to Fig.4).**
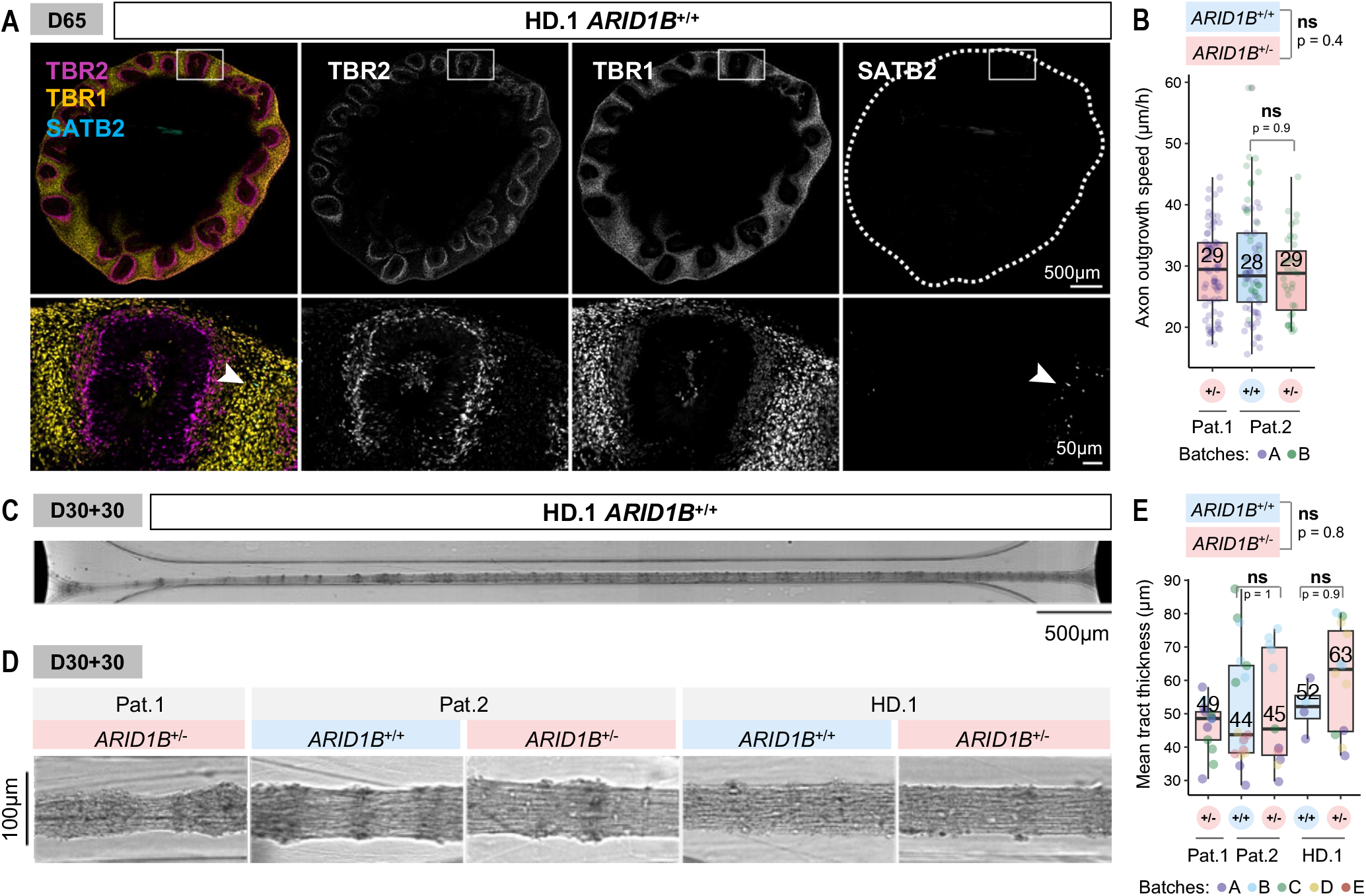
The thickness of axon tracts from early-stage organoids is not affected by *ARID1B* mutations. **(A)** Immunostaining analysis of organoids at D65, showing dorsal cortical intermediate progenitors (TBR2) and excitatory neuron classes (TBR1 subcerebral projection neurons, SATB2 callosal projection neurons). At this stage, organoids are composed of dorsal cortical progenitors and TBR1^+^ and CTIP2^+^ neurons (see **Fig.S3A**); only very few SATB2^+^ neurons have started to differentiate (inset; arrowheads, ➤). **(B)** Quantification of axon outgrowth speed from analysis of live imaging between t = 24 h and t = 72 h (each datapoint is an individual axon). **(C)** Example of a tract from early-stage organoids at D30+30. **(D)** Representative images showing comparable tract thickness in *ARID1B*^+/+^ and *ARID1B*^+/-^ organoids at D30+30. **(E)** Quantification of tract thickness at D30+30 (each datapoint is an individual tract). In boxplots, median values are indicated; each datapoint is an individual axon/tract (technical replicate); datapoint colors indicate organoid batches (biological replicates). Statistical tests are analysis of variance (ANOVA); 0 ≤ p < 0.001, ***; 0.001 ≤ p < 0.01, **; 0.01 ≤ p < 0.05, *; p ≥ 0.05, ns (see results of statistical tests in **Table S4**).

**Table S1.**
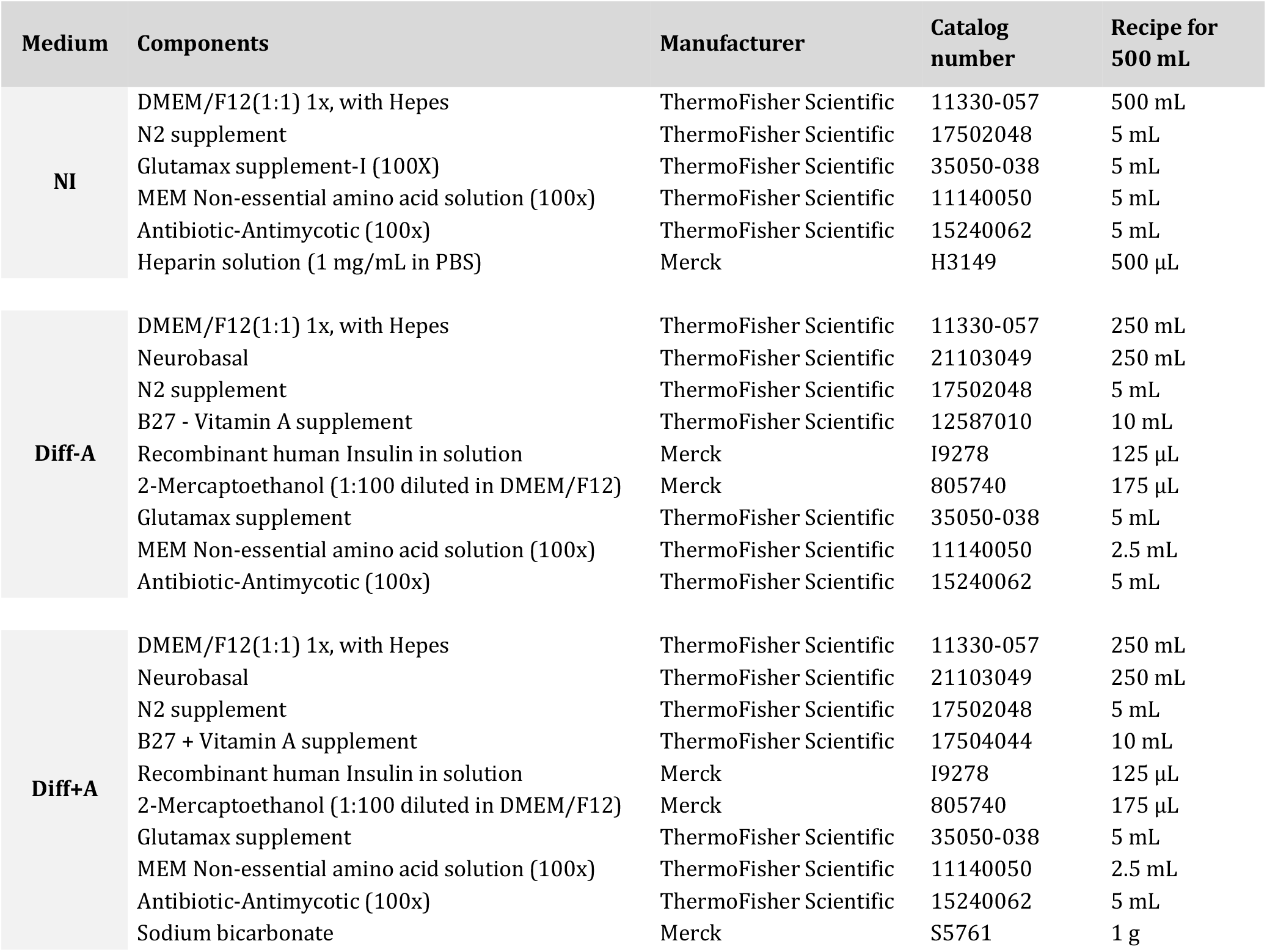
Medium composition for telencephalic organoid culture. Medium components and recipe for 500 mL of Neural Induction medium (NI), Differentiation medium without vitamin A (Diff-A), and Differentiation medium with Vitamin A (Diff+A).

| Medium | Components | Manufacturer | Catalog number | Recipe for 500 mL |
| --- | --- | --- | --- | --- |
| NI | DMEM/F12(1:1) 1x, with Hepes | ThermoFisher Scientific | 11330-057 | 500 mL |
|  | N2 supplement | ThermoFisher Scientific | 17502048 | 5 mL |
|  | Glutamax supplement-I (100X) | ThermoFisher Scientific | 35050-038 | 5 mL |
|  | MEM Non-essential amino acid solution (100x) | ThermoFisher Scientific | 11140050 | 5 mL |
|  | Antibiotic-Antimycotic (100x) | ThermoFisher Scientific | 15240062 | 5 mL |
|  | Heparin solution (1 mg/mL in PBS) | Merck | H3149 | 500 µL |
| Diff-A | DMEM/F12(1:1) 1x, with Hepes | ThermoFisher Scientific | 11330-057 | 250 mL |
|  | Neurobasal | ThermoFisher Scientific | 21103049 | 250 mL |
|  | N2 supplement | ThermoFisher Scientific | 17502048 | 5 mL |
|  | B27 - Vitamin A supplement | ThermoFisher Scientific | 12587010 | 10 mL |
|  | Recombinant human Insulin in solution | Merck | I9278 | 125 µL |
|  | 2-Mercaptoethanol (1:100 diluted in DMEM/F12) | Merck | 805740 | 175 µL |
|  | Glutamax supplement | ThermoFisher Scientific | 35050-038 | 5 mL |
|  | MEM Non-essential amino acid solution (100x) | ThermoFisher Scientific | 11140050 | 2.5 mL |
|  | Antibiotic-Antimycotic (100x) | ThermoFisher Scientific | 15240062 | 5 mL |
| Diff+A | DMEM/F12(1:1) 1x, with Hepes | ThermoFisher Scientific | 11330-057 | 250 mL |
|  | Neurobasal | ThermoFisher Scientific | 21103049 | 250 mL |
|  | N2 supplement | ThermoFisher Scientific | 17502048 | 5 mL |
|  | B27 + Vitamin A supplement | ThermoFisher Scientific | 17504044 | 10 mL |
|  | Recombinant human Insulin in solution | Merck | I9278 | 125 µL |
|  | 2-Mercaptoethanol (1:100 diluted in DMEM/F12) | Merck | 805740 | 175 µL |
|  | Glutamax supplement | ThermoFisher Scientific | 35050-038 | 5 mL |
|  | MEM Non-essential amino acid solution (100x) | ThermoFisher Scientific | 11140050 | 2.5 mL |
|  | Antibiotic-Antimycotic (100x) | ThermoFisher Scientific | 15240062 | 5 mL |
|  | Sodium bicarbonate | Merck | S5761 | 1 g |

**Table S2.** Antibodies used in this study. IF, immunofluorescence; WB, western blotting; FC, flow cytometry.

| Primary antibodies |  |  |  |  |
| --- | --- | --- | --- | --- |
| Species | Antigen | Manufacturer | Catalog# | Application |
| Mouse | SATB2 | Abcam | ab51502 | IF |
| Ms | ARID1B | Santa Cruz Biotechnology | sc-32762 | IF |
| Rb | ARID1A | Abcam | ab182560 | IF |
| Rabbit | SMARCC1/BAF155 | Cell Signaling Technology | 11956S | IF |
| Goat | SOX2 | R&D Systems | AF2018 | IF |
| Sheep | TBR2 | R&D Systems | AF6166 | IF |
| Rabbit | TBR1 | Abcam | ab31940 | IF |
| Rat | CTIP2 | Abcam | ab18465 | IF |
| Rb | dsRed | Clontech | 632496 | IF |
| Ms | ARID1A | Abcam | ab242377 | WB |
| Rb | ARID1B | Cell Signalling Technology | 92964 | WB |
| Rb | BAF155 | Cell Signalling Technology | 11956S | WB |
| Rb | BAF170 | Thermo Scientific | PA5-54351 | WB |
| Rb | b-Actin | Abcam | ab8227 | WB |

| Secondary antibodies |  |  |  |
| --- | --- | --- | --- |
| Antibody | Manufacturer | Catalog# | Application |
| Alexa Fluor 488 Donkey anti-rabbit | Invitrogen | A21206 | IF |
| Alexa Fluor 488 Donkey anti-mouse | Invitrogen | A21202 | IF |
| Alexa Fluor 488 Donkey anti-chicken | Jackson ImmunoResearch | 703-545-155 | IF |
| Alexa Fluor 488 Donkey anti-goat | Invitrogen | A11055 | IF |
| Alexa Fluor 568 Donkey anti-rabbit | Invitrogen | A10042 | IF |
| Alexa Fluor 568 Donkey anti-mouse | Invitrogen | A10037 | IF |
| Alexa Fluor 568 Donkey anti-goat | Invitrogen | A11057 | IF |
| Alexa Fluor 647 Donkey anti-rabbit | Invitrogen | A31573 | IF |
| Alexa Fluor 647 Donkey anti-mouse | Invitrogen | A31571 | IF |
| Alexa Fluor 647 Donkey anti-goat | Invitrogen | A21447 | IF |
| Alexa Fluor 647 Donkey anti-rat | Jackson ImmunoResearch | 712-605-150 | IF |
| IRDye® 800CW Goat anti-Mouse IgG | LI-COR Biosciences | 926-32210 | WB |
| IRDye® 680RD Goat anti-Rabbit IgG | LI-COR Biosciences | 926-68071 | WB |

| Conjugated antibodies |  |  |  |
| --- | --- | --- | --- |
| Antibody | Manufacturer | Catalog# | Application |
| Alexa Fluor® 488 Mouse anti-Human TRA-1-60 (Clone TRA-1-60) | BD Biosciences | 560796 | FC |
| Alexa Fluor® 647 Mouse anti-SSEA-4 (Clone MC813-70) | BD Biosciences | 560173 | FC |

**Table S3.** Lysis buffer for protein sample preparation.

| Solution | Components | Recipe for 1 L |
| --- | --- | --- |
| Protein lysis buffer | **RIPA buffer (store at 4 °C):** |  |
|  | 20mM Tris-HCl (pH 7.5) | 20 mL from 1M Stock |
|  | 150mM NaCl | 30 mL from 5M Stock |
|  | 1mM Na<sub>2</sub>EDTA | 2 mL from 500M Stock |
|  | 1mM EGTA | 0,38 g (MW 380,35 g/mol) |
|  | 1% NP-40 | 10 mL |
|  | 0.1% SDS | 10 mL 10% Stock solution |
|  | MilliQ H<sub>2</sub>O | 927 mL |
|  | **Add freshly before use:** |  |
|  | PhosStop (Roche, 4906837001) | 1 tablet / 10mL of RIPA |
|  | Complete Mini EDTA free protease inhibitor (Roche, 11836170001) | 1 tablet / 10mL of RIPA |

**Table S4.** Results of statistical analyses. Analysis of variance (ANOVA); 0 ≤ p < 0.001, ***; 0.001 ≤ p < 0.01, **; 0.01 ≤ p < 0.05, *; p ≥ 0.05, ns. Whenever possible, the influence on the quantitative variable was calculated from the interaction between *ARID1B* genotype (WT-mut) and genetic background (Pat.1, Pat.2, and HD.1), to obtain isogenic comparisons (WT:Pat2-mut:Pat2 and WT:HD.1-mut:HD.1). When there were not enough observations per each genetic background, only the *ARID1B* genotype was considered (WT-mut). WT, *ARID1B*^+/+^; mut, *ARID1B*^+/-^.

| Figure: data | Comparison | Mean difference | Lower range | Upper range | Adjusted p-value |  |
| --- | --- | --- | --- | --- | --- | --- |
| Fig.S2C: WB, iPSCs, ARID1B | WT-mut | -0.4375 | -0.84932 | -0.02568 | 0.043062 | * |
| Fig.S3C: WB, OrgD60, ARID1B | WT-mut | -0.625 | -1.17994 | -0.07006 | 0.033983 | * |
| Fig.S3D: WB, OrgD60, ARID1A | WT-mut | -0.19 | -1.2453 | 0.865296 | 0.662939 | ns |
| Fig.S3E: WB, OrgD60, BAF155 | WT-mut | -0.29 | -0.91953 | 0.33953 | 0.289567 | ns |
| Fig.S3F: WB, OrgD60, BAF170 | WT-mut | 0.141667 | -0.20469 | 0.48802 | 0.3412 | ns |
| Fig.S4A: WB, OrgD120, ARID1B | WT-mut | -0.46194 | -0.64051 | -0.28338 | 4.25E-05 | *** |
| Fig.S4D: SATB2<sup>+</sup> neurons/mm<sup>2</sup> | WT-mut | 29.77637 | -25.7557 | 85.30846 | 0.284986 | ns |
|  | WT:Pat2-mut:Pat2 | 34.8056 | -98.3449 | 167.9562 | 0.968993 | ns |
|  | WT:HD.1-mut:HD.1 | 75.6754 | -45.7655 | 197.1163 | 0.438014 | ns |
| Fig.S5: %Cell/cluster (excitatory lineage) | IPCs: WT-mut | 3.334026 | 0.684653 | 5.983399 | 0.016223 | * |
|  | ExN1: WT-mut | 0.155714 | -2.09261 | 2.404039 | 0.886576 | ns |
|  | ExN2: WT-mut | -3.08734 | -5.20272 | -0.97196 | 0.006402 | ** |
|  | ExN3: WT-mut | -0.40312 | -2.83466 | 2.028425 | 0.733086 | ns |
| Fig.3F: SATB2<sup>+</sup>/Crimson<sup>+</sup> cells (%) | WT-mut | -5.10486 | -14.4889 | 4.27923 | 0.277714 | ns |
|  | WT:Pat2-mut:Pat2 | -3.12599 | -24.6068 | 18.35485 | 0.99783 | ns |
|  | WT:HD.1-mut:HD.1 | -3.46927 | -26.2911 | 19.35252 | 0.99733 | ns |
| Fig.3G: Crimson<sup>+</sup> cells per 100k cells | WT-mut | -37.2031 | -49.8242 | -24.5819 | 1.28E-07 | *** |
|  | WT:Pat2-mut:Pat2 | -36.9688 | -63.5534 | -10.3842 | 0.001616 | ** |
|  | WT:HD.1-mut:HD.1 | -22.9974 | -56.1671 | 10.17224 | 0.335426 | ns |
| Fig.3H: Mean tract thickness (D120 tracts) | WT-mut | 14.38672 | 11.10506 | 17.66837 | 2.55E-15 | *** |
|  | WT:Pat2-mut:Pat2 | 10.01818 | 3.412554 | 16.6238 | 0.000304 | *** |
|  | WT:HD.1-mut:HD.1 | 11.64495 | 3.322536 | 19.96737 | 0.001148 | ** |
| Fig.3I: % Axon-associated cells | WT-mut | -11.1215 | -14.0463 | -8.19665 | 7.85E-09 | *** |
|  | WT:Pat2-mut:Pat2 | -10.7147 | -17.7935 | -3.63594 | 0.000865 | *** |
|  | WT:HD.1-mut:HD.1 | -9.0855 | -15.8692 | -2.3018 | 0.003715 | ** |
| Fig.4D: Mean number of axons (0 to 160um) | WT-mut | -15.2435 | -21.2193 | -9.26773 | 8.64E-07 | *** |
|  | WT:Pat2-mut:Pat2 | -13.1394 | -26.1532 | -0.12567 | 0.0463 | * |
|  | WT:HD.1-mut:HD.1 | -15.4418 | -29.9166 | -0.96705 | 0.028853 | * |
| Fig.4F: Axon outgrowth speed (D120 tracts) | WT-mut | 1.882407 | 0.227716 | 3.537097 | 0.025825 | * |
|  | WT:Pat2-mut:Pat2 | 0.984848 | -2.57987 | 4.54957 | 0.969314 | ns |
|  | WT:HD.1-mut:HD.1 | 1.041021 | -2.79772 | 4.879763 | 0.971695 | ns |
| Fig.S11B: Axon outgrowth speed (D30 tracts) | WT-mut | 1.01212 | -1.31642 | 3.340655 | 0.392125 | ns |
|  | WT:Pat2-mut:Pat2 | -0.61484 | -4.36372 | 3.134044 | 0.920523 | ns |
| Fig.S11C: Mean tract thickness (D30 tracts) | WT-mut | -1.07344 | -9.38344 | 7.236555 | 0.796498 | ns |
|  | WT:Pat2-mut:Pat2 | -0.51505 | -17.7813 | 16.75119 | 0.999999 | ns |
|  | WT:HD.1-mut:HD.1 | -8.72619 | -34.4882 | 17.03577 | 0.915146 | ns |

